# Rictor CRISPR Gene Editing by Lipid Nanoparticle Delivery Stimulates Anti-tumor Immunity in Breast Cancer Liver Metastasis Model

**DOI:** 10.1101/2025.06.26.661471

**Authors:** Yaqoob Ali, Tyler Galbraith, Nourhan Abdelfattah, Arturas Ziemys, Thomas L Wong, Chihiro Hashimoto, Maryam Faisal, Xu Qian, Harlan Cook, Tej Pandita, Yitian Xu, Roberto Rosato, Shu-hsia Chen, Kyuson Yun, Fransisca Leonard

## Abstract

Triple-negative breast cancer (TNBC) is a highly aggressive subtype, accounting for 10–15% of breast cancer cases in the United States. Liver metastases, common in advanced TNBC, are linked to especially poor outcomes, with a 5-year survival rate of just 11%. Although immune checkpoint inhibitors (ICIs) targeting PD-1 or PD-L1 show promise, durable responses in TNBC remain uncommon. This is largely due to a profoundly immunosuppressive tumor microenvironment (TME), driven by tumor-associated myeloid cells. Tumor-associated macrophages (TAMs) and neutrophils (TANs) polarize into immunosuppressive M2 and N2 phenotypes, respectively, suppressing T cell activity through cytokines, ROS, and checkpoint ligands such as VISTA. Myeloid-derived suppressor cells (MDSCs) further inhibit immunity by depleting nutrients and inducing regulatory T cells. As a result, despite its immunogenic features, TNBC remains resistant to immunotherapy due to persistent myeloid-mediated suppression.

Here, we developed ionizable lipid nanoparticles (iLNPs) engineered to deliver the CRISPR-Cas12a ribonuclease complex targeting Rictor, a critical component of mTORC2, for *in vivo* reprograming of myeloid cells. The intravenous (IV) injection of CRISPR Rictor-targeting iLNP (CR-Ric-LNP) showed efficient uptake by circulating myeloid cells and accumulation into the breast cancer liver metastases. Notably, Rictor gene editing triggered pro-inflammatory activation of myeloid cells in the TME, enhancing antitumor responses. Single-cell RNA sequencing revealed that *Rictor* silencing treated samples showed induced rapid remodeling of the TME, with a significant reduction in immunosuppressive macrophages within 24 hours of treatment. Concurrently, cytotoxic T-cell populations exhibited increased interferon-gamma (*Ifng*) production, driving the emergence of specific myeloid clusters that were responsive to Interferon signaling, particularly in macrophages and neutrophils. A shift from an immunosuppressive to an inflammatory TME was further evidenced by an elevated *Cxcl10/Spp1* ratio in myeloid cells. CR-Ric-LNP treatment also enhanced T-cell activation, reducing exhausted T cells and regulatory T cells (Tregs) while expanding natural killer (NK) cells, naïve CD4+, and CD8+ T cells. These changes correlated with a decreased proportion of tumor cells and proliferating cells, ultimately leading to a significant survival benefit in a 4T1 breast cancer liver metastasis model. Our findings demonstrate that myeloid-targeted *Rictor* silencing reprograms the TME, promoting antitumor immunity and improving therapeutic outcomes.

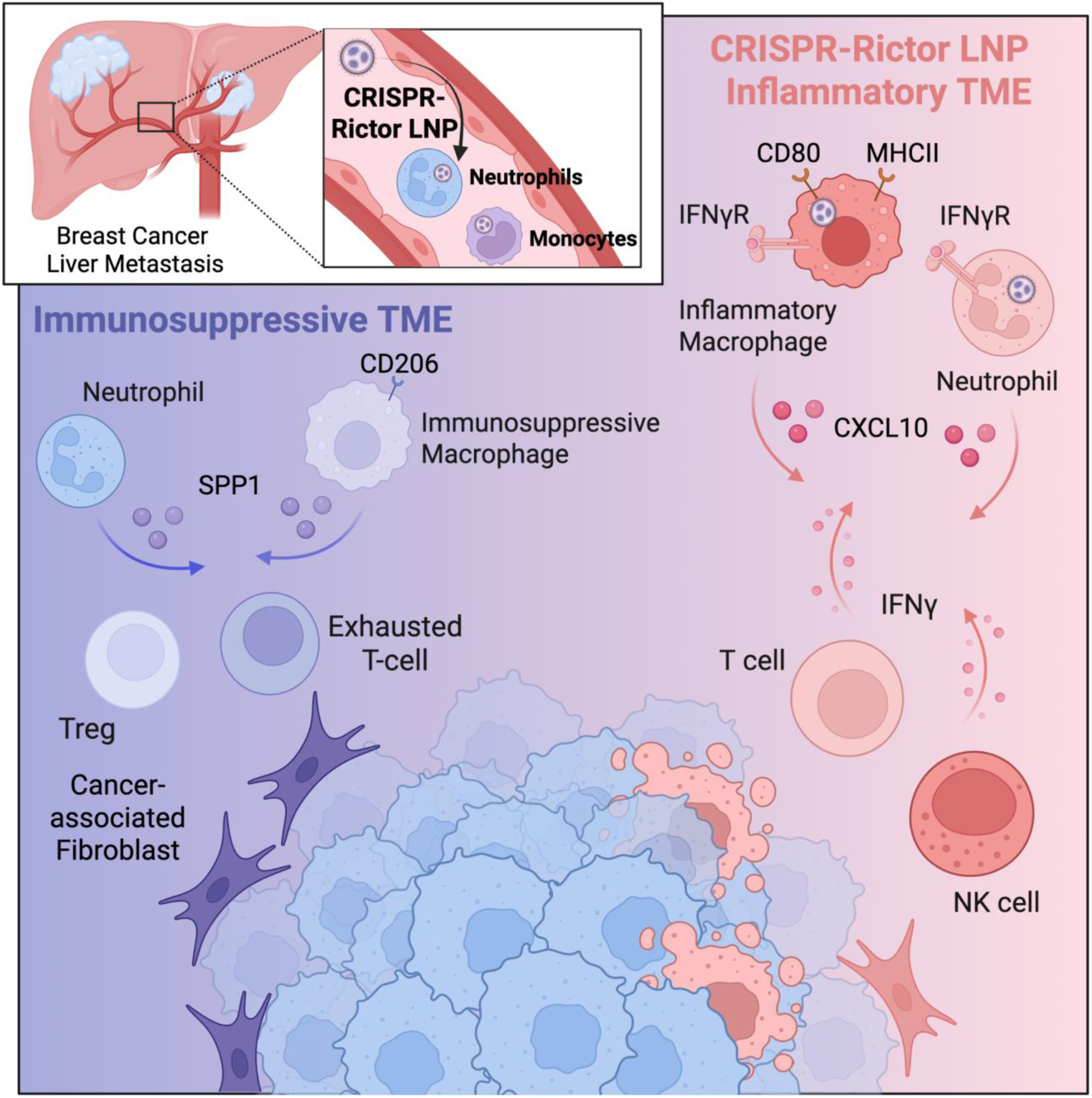

## 1. Introduction

Breast cancer liver metastasis (BCLM) is responsible for poor overall prognosis, with a five-year^1^ survival rate of 11%^2,3^. Current options for treatment often result in treatment resistance and cancer relapse^4–7^, in part due to the immunosuppressive immune system and the TME that play important roles in modulating therapeutic responses in TNBC-LM responses^8,9^.

Immune checkpoint inhibitor (ICI) therapies have revolutionized cancer treatment^10–15^. By inducing antitumor immunity, ICI reactivates cytotoxic cells and memory responses for long-term protection, offering durable, targeted treatment compared to non-specific chemotherapy. ICIs have transformed outcomes for many cancer patients, but their efficacy is still limited to specific subsets of patients with “immune hot” cancers, characterized by an abundance of infiltrating cytotoxic cells that is likely to trigger a strong immune response^16,17^. Current strategies focus on enhancing ICI responsiveness by switching the immune excluded “immune cold” cancers with low cytotoxic cell infiltration into “immune hot” cancers^18,19^.

Although immune checkpoint inhibitors (ICIs) targeting PD-1 or PD-L1 have shown promise, sustained responses in TNBC remain rare. This limited efficacy is largely due to the immunosuppressive tumor microenvironment shaped by tumor-associated myeloid cells. Recent studies have revealed that myeloid cells comprising of myeloid-derived suppressor cells (MDSCs), tumor-associate macrophages, and tumor-associated neutrophils are key sub-populations that modulate the immune response in TME^20–23^. These populations are attractive targets for therapy, as their reprograming can enhance ICI response^24,25^. Among the myeloid population, Tumor-associated macrophages (TAMs) are described to exist on a spectrum from pro-inflammatory (M1-like) activity to immunosuppressive or protumor (M2-like) state^26^. TAMs, which may comprise over 50% of the TME, are frequently polarized toward an M2-like phenotype under the influence of IL-10, TGF-β, and STAT3 signaling^27^. These cells promote immune evasion by secreting suppressive cytokines and expressing inhibitory checkpoints such as VISTA, impairing cytotoxic CD8+ T cell function^23^. Likewise, tumor-associated neutrophils (TANs) can adopt an N2 phenotype driven by TGF-β and CXCL8/CXCR2 signaling, which contributes to T cell suppression through reactive oxygen species (ROS) and arginase-1 release^28–32^. In contrast, N1 neutrophils exhibit anti-tumor activity by directly killing cancer cells and recruiting T cells into the TME^28^. Myeloid-derived suppressor cells (MDSCs) comprising monocytic MDSC (mMDSC) and polymorphonuclear MDSC (PMN-MDSC/gMDSC) are subsets of immature myeloid cells which further reinforce this suppression by depleting nutrients like arginine and producing IDO, skewing T cells toward regulatory phenotypes^33,34^. Secreted cytokines from MDSCs and TAMs help establish an immunosuppressive milieu in the TME that promotes increased neovascularization, extracellular matrix reorganization, epithelial-mesenchymal transition, intravasation, and extravasation of cancer stem cells, and the establishment of the pre-metastatic niche^26^. As a result, despite TNBC’s immunogenic profile myeloid-driven immunosuppression remains a major barrier to immunotherapy success.

Transforming and reducing immunosuppressive myeloid cell populations towards a pro-inflammatory/anti-tumorigenic niche offers a promising potential to improve cancer therapy outcomes^35–37^ while maintaining fundamental homeostasis roles. However, myeloid cells exhibit a fluid, dynamic, and reversible, cell state profile inducible by factors in their environment. Despite various approaches targeting TAMs^35–48^, efforts to target monocytes for cancer therapy *in vivo* remain limited. Recent strategies have focused on using nanoparticles, such as CCR2 mRNA-loaded LNP to overexpress CCR2 in monocytes^49^, or apoptotic body-encapsulated CpG modified gold–silver nanorods^50^. Other approaches, such as self-assembled mertansine nanoparticles for suppressing lung metastases^51^ or blocking CD47 using antibody^15,52^ still face challenges from limited drug/mRNA half-life^53^ and the highly plastic nature of monocytes and macrophages that cause phenotypic changes in the TME.

Rapamycin Insensitive Companion of mTOR (*Rictor),* is a critical component of the mTOR2 complex that regulates PI3K/AKT signaling by promoting Akt phosphorylation at Serine 437. The mTOR2 complex is essential for energy and glucose homeostasis regulation^54,55^. In cancer cells, RICTOR is known to regulate and have effects on survival, invasion, proliferation, viability^56^, and metabolic wiring^57^. In gastric and lung cancers^58,59^, overexpression of RICTOR is correlated with a poor prognostic factor, inferences response to therapy^60^, and inhibits cancer cell apoptosis, thereby promoting cancer cell growth via the Akt signaling pathway^61^. In breast cancers, *RICTOR* knockdown attenuated HER2-dependent oncogenesis^59^, while *RICTOR* amplification was reported to promote tumor progression in TNBC. In addition to its known function in cancer cells, RICTOR is also required for M2 macrophage polarization^62–65^. RICTOR deletion has been shown to induces M1-like macrophage polarization in infection^63^, nonalcoholic fatty liver disease^66^, tissue injury^67^ and atherosclerosis^56^ contexts.

To effectively shift the polarization of these immunosuppressive tumor associated myeloid cells to a more pro-inflammatory/anti-tumor state, we target myeloid cell Rictor gene via clustered regularly interspaced short palindromic repeats (CRISPR) system and tested this in a Triple Negative Breast Cancer Liver Metastasis model (TNBC-LM). We accomplished in vivo reprogramming of myeloid cell Rictor by delivering CRISPR construct using ionizable lipid nanoparticles (iLNP). Ionizable lipids, with their unique pH-responsive structure, minimizes immune recognition at neutral pH and become cationic in acidic environments (e.g., endosomes) to enable payload release^68–70^. This payload profile has been previously shown to increase LNP safety profiles and have accelerated the success of iLNP-based vaccine and drugs. Further, by integrating efficacy with biocompatibility, iLNPs have unlocked breakthroughs in gene editing, personalized vaccines, RNA therapies, and enabling researchers to address the previously undruggable molecular targets for tailored therapy development.

In this study, we developed an iLNP system encapsulating Cpf1/Cas12a CRISPR ribonuclease complex targeting *Rictor* (CR-Ric-LNP) to reprogram TME from an immunosuppressive to an inflammatory phenotype.

In this study, we evaluated the performance of our RICTOR-targeting CRISPR complex encapsulated within LNP (CR-Ric-LNP) for the *in vivo* engineering of tumor-associated myeloid cells to reprogram the TME. We have examined the efficacy of the treatment in remodeling TME and eradicating cancer using *in vivo* 4T1 breast cancer liver metastasis mouse model and *in vitro* studies on primary macrophages.

## Results

### CR-LNP design and characterization

We have designed an in-house optimized and validated ionizable LNP and CRISPR construct consisting of Cpf1/Cas12a nuclease and CRISPR guide RNA (crRNA) as a ribonucleoprotein complex (CR-LNP) (Fig 1a) which allow for fast-acting genetic modification without incorporation of genetic material into host cells. Cas12a is a CRISPR nuclease with higher fidelity compared to Cas9 ^71,72^. This highly versatile system can be used to target other genes by swapping the specific guide RNA sequences (Fig 1a). Ionizable lipid DLin-MC3-DMA (pKa=6.44) in addition to phosphatidylcholine and cholesterol was utilized to form iLNP for loading CRISPR system at pH of 5.5. Varying concentration of lipid and CRISPR nuclease/crRNA compositions were tested on bone marrow-derived macrophages (BMDMs) to observe the effects of the CRISPR-LNP on viability. The ratio between CRISPR nuclease and crRNA was kept to 1:1 to enable optimal complex formation of CRISPR system. Our results indicated that the increase in CRISPR content did not cause any toxicity to BMDMs (Fig 1b). Lipid concentration up to 4mg/mL did not affect macrophage viability, while concentration of 10mg/mL lipid led to ∼25% toxicity to BMDMs (Fig 1b). We thus formulated the CRISPR-Liposome using 4mg/mL lipid concentration and 80µg crRNA content, which resulted in LNP with the size of 167.4±2.7 nm, average polydispersity index of 0.08, surface zeta potential of around -7.3±1.1 mV, and CRISPR encapsulation efficiency of >77% consistently (Fig 1c). We initially tested the uptake of rhodamine-tagged CR-LNP in Thp-1 cells, a monocytic cell line. LNP uptake was observed at 1h after treatment (Fig 1d). We evaluated the activity of CRISPR-loaded lipid nanoparticles (CR-LNPs) in multiple cancer cell lines and primary bone marrow-derived macrophages. Initially, GFP-expressing MDA-MB-463 cells were treated with GFP-targeting CR-LNPs, resulting in a significant reduction in GFP fluorescence after 72 hours (Suppl Fig 1a). DNA cleavage upon CRISPR treatment with CR-GFP-LNP in the MDA-MB-463 was validated using the GeneArt™ Genomic Cleavage Detection Kit, demonstrating ∼30% editing efficiency across three independent replicates, as evidenced by the presence of cleaved DNA fragments compared to the untreated control (Suppl Fig 1b).

**Figure 1.**
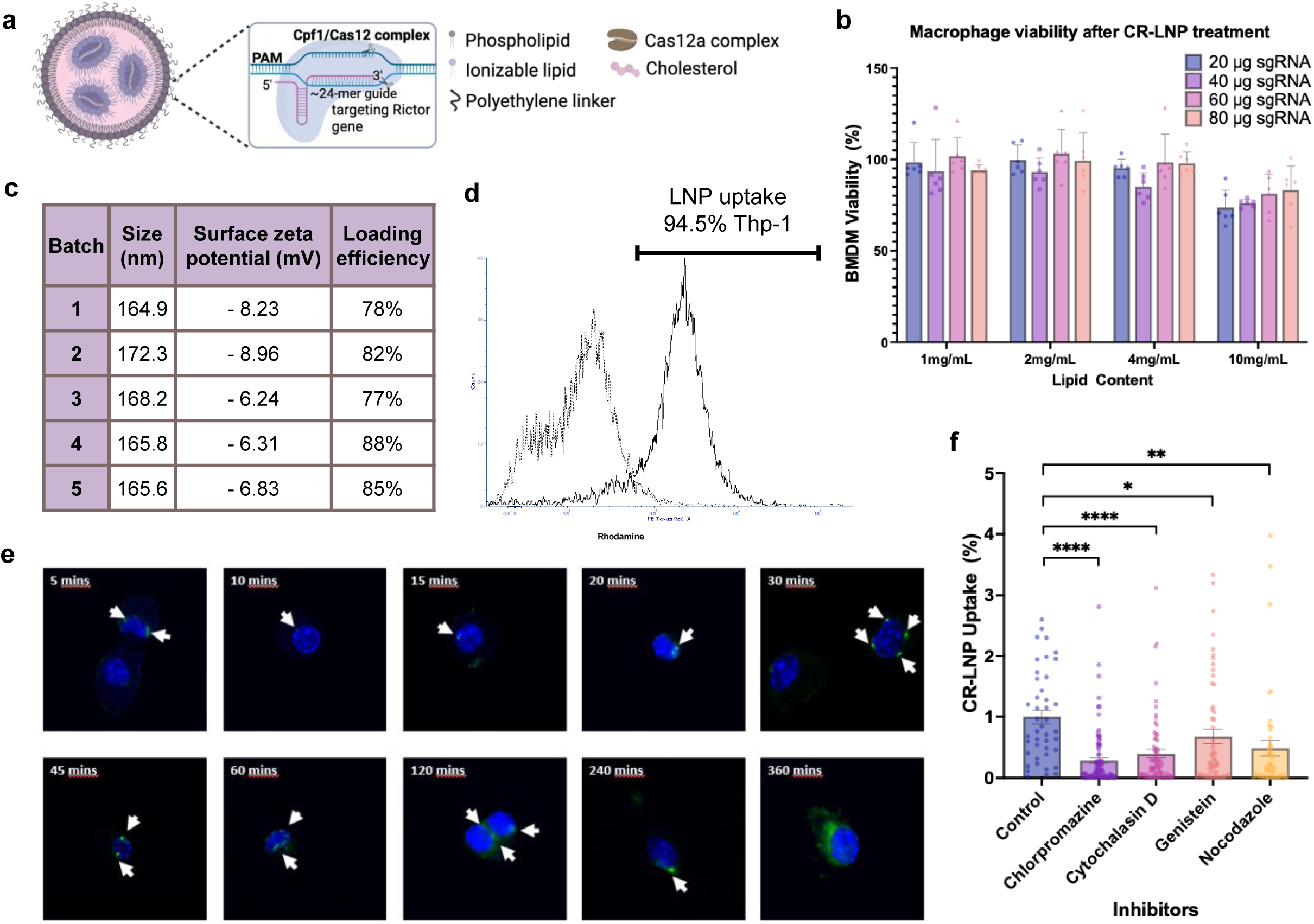
CR-LNP optimization and characterization. a) Schematic of CR-LNP, composition and encapsulation of CRISPR component. b) Varying LNP compositions tested for cytotoxicity towards macrophages. c) Size and zeta Potential of CR-LNP characterized by dynamic light scattering. d) Flow cytometry analysis of LNP uptake by Thp1 2h after treatment. e) Representative immunofluorescence images of γ-H2AX foci (green) indicating double strand breaks in primary murine macrophages following CRISPR-mediated DNA targeting. Nuclei were counterstained with DAPI (blue). f) Uptake inhibitor study to evaluate CR-LNP uptake mechanism. Primary bone marrow derived macrophages were treated with uptake inhibitors for 4 hours before incubation with CR-LNPs. LNP uptake was measured 2 hours after incubation. Data presented as cellular uptake normalized to mean CR-LNP uptake in control untreated macrophages. Mean±SEM.

To determine the relationship between cellular uptake and editing efficiency, RhodamineB-conjugated luciferase-targeting CR-LNPs were applied to two cancer cell lines with varying CR-LNP uptake rates: 4T1 (breast cancer) and BxPC3 (pancreatic cancer). Fluorescence analysis demonstrated nearly 2-fold higher CR-LNP uptake in 4T1 cells compared to BxPC3 (Suppl Fig 1c). Consistent with this observation, luciferase knockdown efficiency was significantly greater in 4T1 cells (∼50% reduction) than in BxPC3 cells (∼20% reduction) (Suppl Fig 1c).

In primary murine macrophages, CRISPR-mediated DNA double strand break (DSB) was confirmed by the formation of γ-H2AX foci, detectable as early as 5 minutes post-treatment and persisting for up to 4 hours (Fig 1e). We also evaluated the effects of various pharmacological inhibitors implicated in various cellular uptake mechanisms to assess the uptake mechanism of the CR-LNP. Among the tested compounds, chlorpromazine demonstrated the highest inhibitory activity, achieving nearly 75% inhibition of uptake (Fig 1f). Cytochalasin D also exhibited significant inhibition, reducing uptake by 60%, while nocodazole caused a moderate reduction of 50%. In contrast, genistein showed the weakest effect, with only ∼30% inhibition observed. These findings suggest clathrin-mediated endocytosis as a major uptake pathway for CR-LNP. The intermediate effects of cytochalasin D and nocodazole indicate partial dependence on actin dynamics and microtubule integrity, respectively, whereas the modest inhibition by genistein implies a lesser role for tyrosine kinase-mediated mechanisms in the process.

### Screening for candidate genes and *in vitro* assessments

To identify candidate genes that can reprogram immunosuppressive macrophage polarization, we conducted functional screening in murine BMDM. From an extensive literature review, we identified and evaluated a panel of 20 different genes reported to regulate M1/M2 polarization such as various cytokine genes, genes in Arginase pathways^73^, PI3K^74^, and mTOR pathways^63,73^. We treated murine BMDMs with CR-LNP targeting candidate genes, analyzed CD80 (M1 marker) and CD206 (M2 marker) expression after 1 day of treatment as functional markers of macrophage polarization using immunofluorescence (IF) staining and fluorescence microscopy analysis. Single-gene disruption of Rictor induced functional repolarization of M2-conditioned macrophages, as evidenced by enhanced CD80 expression (Fig 2a). Despite continuous M2-polarizing conditions (addition of conditioned media from 4T1 breast cancer cell line), genetic perturbation of *Rictor* via CR-Ric-LNP elicited a marked phenotypic shift in macrophages, characterized by significant increase of CD80 expression (Figure 2b).

**Figure 2.**
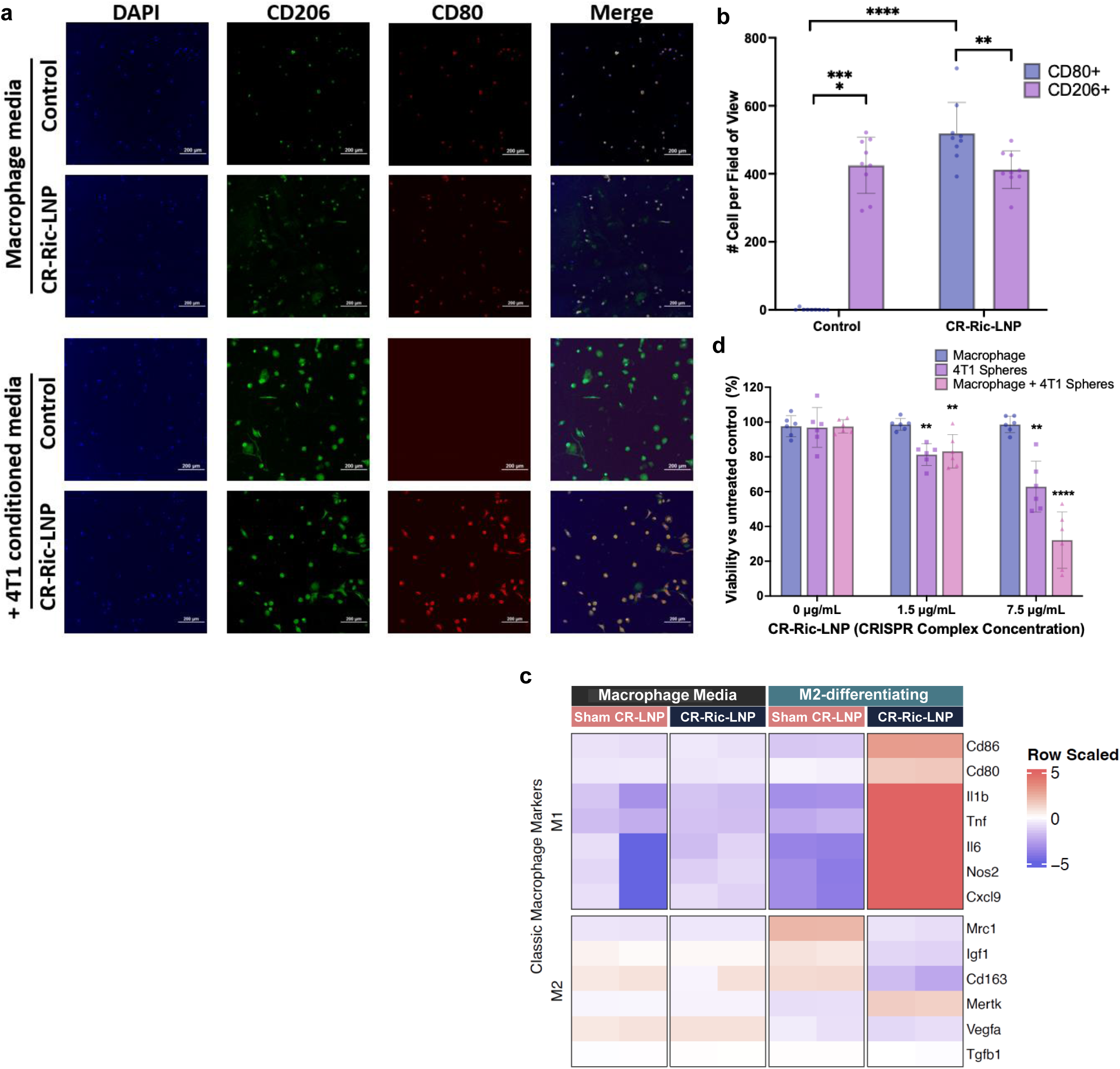
CRISPR screening targeting _c_ immunosuppressive genes using CR-LNP platform. a) Representative immunofluorescence staining of bone marrow derived murine macrophages treated with CR-LNP targeting various immunosuppressive macrophage genes. Staining to detect inflammatory macrophage marker (CD80) and immunosuppressive macrophage marker (CD206). b) Quantification of number of cells expressing CD206 and CD80 after various CRISPR treatments. Data are displayed as mean ± SD, n=9, **p<0.01, ****p<0.0001. c) Gene expression data from RNA sequencing data of CR-Ric-LNP vs sham CR-LNP (CR-Cont) under the presence of M1 (LPS+IFNƳ) or M2 (IL-4+M-CSF) growth factors. d) Viability assessment of macrophages and 4T1 tumor spheres in single and coculture as response to CR-RIC-LNP treatment. Measurement was conducted 24h after treatment and data are displayed as mean ± SD, n=6. **P<0.01, ****P<0.0001 to untreated control.

The genomic sequencing of *Rictor* exon targeted by CR-Ric-LNP revealed numerous mutations in macrophages, with a representative mutation profile (Suppl Fig 1d).

Bulk RNA sequencing data from murine BMDM revealed that knocking out *Rictor* reduced the expression of multiple M2 markers (*Mrc1, CD163, and Igf1*) and increased several M1 markers (*Cd86, Cd80, Il-1β, Tnfα, Il-6, and Nos2*) at the RNA level (Fig. 2c)^62^.

We next assessed the effects of *Rictor* gene editing on both macrophages and 4T1 cancer cells. We have previously shown that cell monolayer tend to overestimate drug efficacy and less relevant for clinical translation, since spheres are harder to penetrate and can be up to 30x more resistant for drug targeting^75^. In our single and coculture studies, CR-Ric-LNP treatment reduced 4T1 viability, and the effect was more pronounced in the presence of macrophages, indicating potential synergy from the dual targeting of this treatment (Fig 2d). Further, when 4T1 spheres were cocultured together with a mixture of CR-Ric-LNP-treated and untreated M2-differentiated macrophages, there was a greater than 3-fold increase in macrophage infiltration compared to control, with majority of infiltrating macrophages expressing CD80 (Suppl Fig 1f).

### CR-Ric-LNP are taken up by myeloid cells and accumulate in breast cancer liver metastases

To determine the cell type taking up the LNP in circulation, we incubated the murine peripheral blood mononuclear cells with rhodamine-tagged CR-Ric-LNP for over 2 hours. More than 12% of immune cells internalized CR-LNPs (Fig 3a), with >95% taken up by CD11b⁺ myeloid cells (Fig 3b). Further subset analysis based on Ly6G and Ly6C expression revealed that Ly6G⁺ neutrophil/gMDSCs accounted for the majority of CR-LNP uptake (Fig. 3c). A time-dependent increase in particle uptake was observed between 10 minutes and 120 minutes, although this trend did not reach statistical significance (Fig 3d). Ly6G⁺ cells (e.g., neutrophils) accounted for 80– 93% of total myeloid cells, whereas Ly6G⁻ monocytic cells represented only 3–5%. Despite this disparity in population size, uptake rates between Ly6G⁺ and Ly6G⁻ cells were comparable, up to around 12% and 15% in granulocytic and monocytic lineages after 2 hours, respectively (Fig 3d).

**Figure 3.**
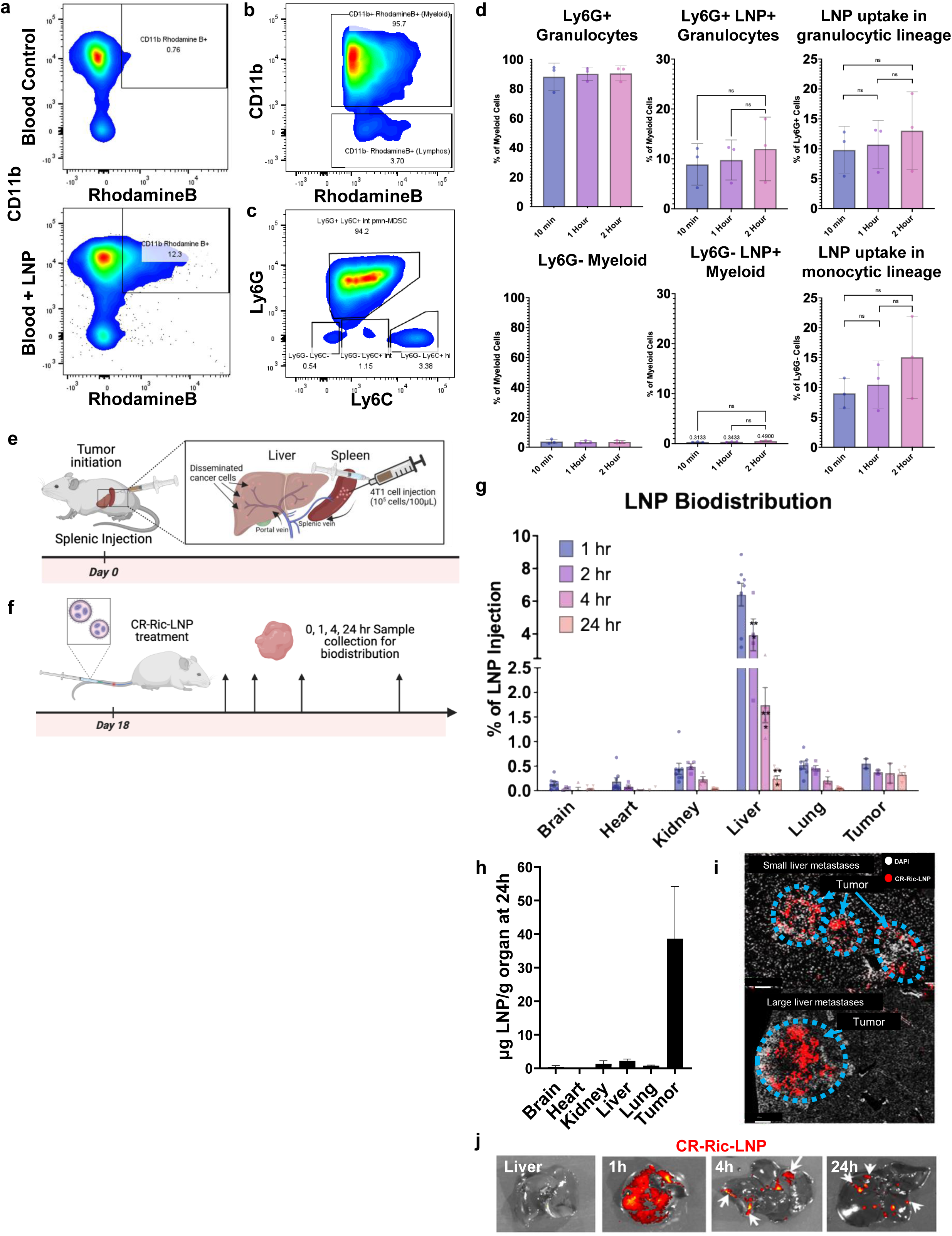
Biodistribution and efficacy of rhodamine-tagged CR CR-RIC-LNP in whole blood and liver metastases. a) Detection of LNP uptake by CD11b+ cells in the whole blood, gated from CD45+ cells. b) Representative scatter plot of CD45+ RhodamineB+ gated population showed predominant uptake of the LNP by CD11b+ myeloid cells. c) Gated CD11b+ RhodamineB+ plotted against Ly6C and Ly6G showed predominant uptake by Ly6Ghi and secondary uptake by Ly6G-Ly6Chi populations. d) Quantification of particle uptake from scatterplots (c), Ly6G+ and Ly6G-myeloid cell composition, as well as normalized uptake by both populations. Biodistribution was observed by IVIS and fluorescent imaging of the liver (e,f,g,h). e) Schematic of breast cancer liver metastasis initiation. 4T1 TNBC cells were injected into the spleen, followed by splenectomy. f) Schematic of biodistribution study. Splenic injection surgery for tumor initiation was conducted at D0, and animals were treated with CR-Ric-LNP at D18, when macro-metastases have already formed in the liver. Organs were harvested at 0h, 1h, 4h, and 24h after treatment for biodistribution measurements. g) Kinetic study of CR-Ric-LNP biodistribution as measured by organ homogenates. h) Calculated concentration of CR-LNP in various organs and liver metastatic lesions 24h after treatment. i) High magnification of liver fluorescent imaging detection of rhodamine signal from CR-LNP 24h after treatment confirmed the organ homogenates results. j) Kinetics study of Rhodamine-tagged CR-Ric-LNP showed the wash out of CR-LNP from the liver and retention within metastatic lesions after 4h-24h of treatment (arrowheads).

To track LNP biodistribution in TNBC liver metastases, we utilized the 4T1 liver metastasis model in mice established via splenic injection of 4T1 cells followed by splenectomy (Fig 3e). Eighteen days after metastasis initiation, CR-Ric-LNPs with rhodamine-tagged phospholipids were injected intravenously and were detected via IVIS imaging and fluorescence measurements of organ homogenates (Fig. 3f). Fluorescence measurement of organ homogenates showed that CR-Ric-LNPs were accumulated in the liver within the first hours (Figs. 3g, j). By 24 hours, the accumulated CR-LNP in the liver was cleared, CR-Ric-LNP were enriched in the metastatic lesions, accounting for ∼40 µg/g tissue, equal to ∼25nM) (Fig. 3g, h). These data were consistent with IVIS and fluorescence imaging of the liver organs (Fig 3i, j). These results demonstrated that our approach can overcome a major hurdle in delivery of LNP into cancer: the difficulty to retain nanoparticles within the liver metastases due to the stromal composition and internal pressure of the lesions^75^.

### CR-Ric-LNP treatment in liver metastasis remodels TME and promotes anti-tumor immune response

To gain mechanistic insight how the CR-Ric-LNP treatment affects the TME, we used single cell RNA-sequencing (scRNAseq). To mimic clinical conditions where liver metastasis is typically detected at the macro metastases stage, we administered CR-Ric-LNP at day 18 post-initiation of liver metastases (Fig 4a), a time point at which most 4T1 mice had developed macro metastases of >1mm diameter in the liver. Mice were harvested at 24h post treatment and metastatic lesions were pooled from 3 animals each from control and CR-Ric-LNP treated animals to increase throughput and mitigate individual variations.

**Figure 4.**
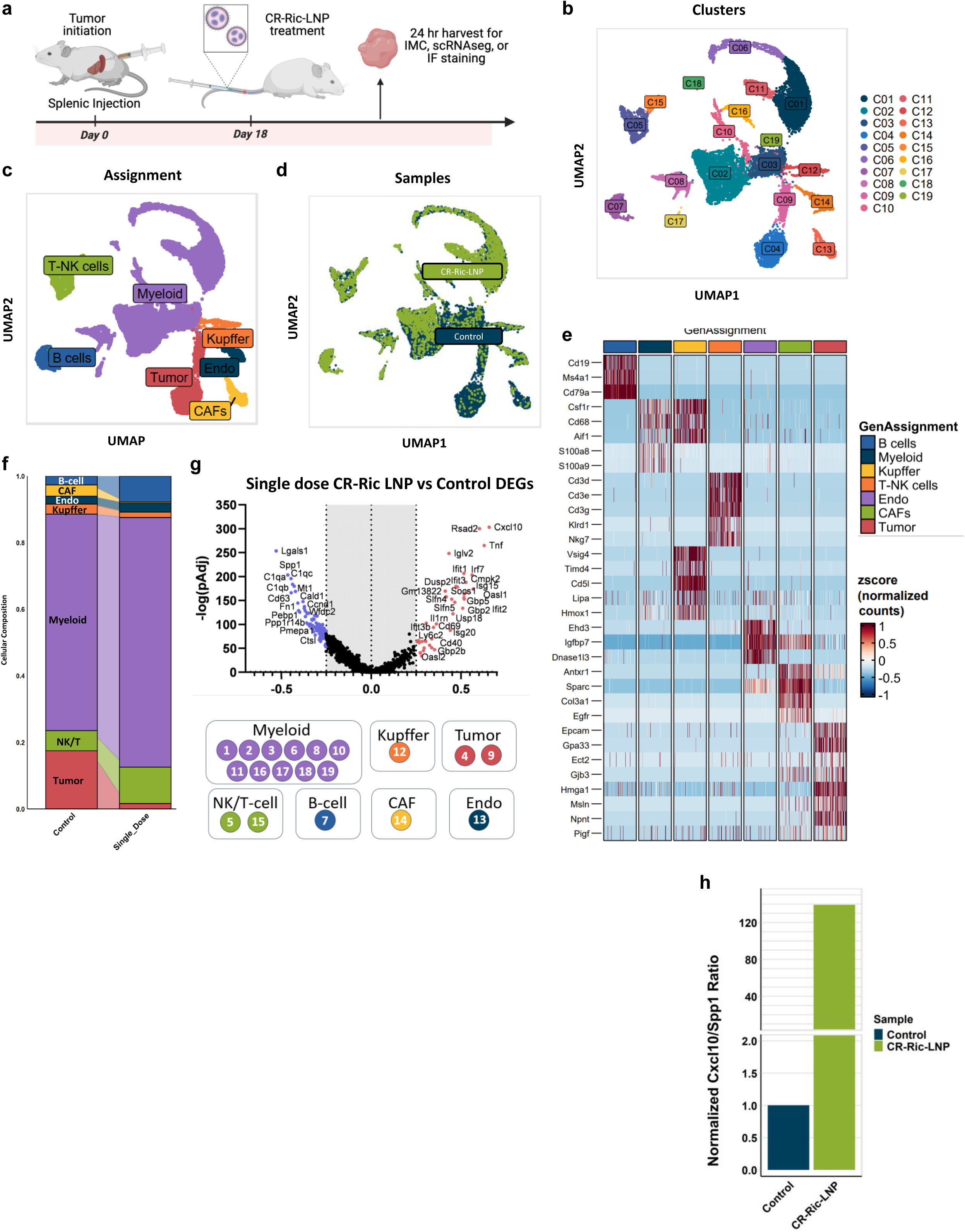
scRNAseq analysis of 4T1 murine model of liver metastasis after treatment with CR-Ric-LNP. a) Schematic of in *vivo* experimental design b) UMAP projection and unsupervised clustering of the liver metastasis TME. c) UMAP of cell assignment based on known gene markers. d) UMAP projection of single cells from control and Rictor-KD liver mets cells. e) Heatmap indicating normalized expression of top DEG in each cell populations. f) Normalized quantitation showing different cellular composition in control and CR-Rictor-LNP-treated samples. g) Volcano plot of DEG after single dose of CR-Ric-LNP. The logFC indicates the mean expression level for each gene. Each dot represents one gene, red dots represent upregulated genes and blue dots downregulated genes. h) Fold change expression ratio of Cxcl10 to Spp1 across samples in breast cancer liver metastasis represented as barplots.

We analyzed and integrated 21,185 single cells that passed all QC steps from both datasets. Non-linear dimensionality reduction was performed and shown in uniform manifold approximation and projection (UMAP) (Fig 4b). Unsupervised clustering using Seurat workflow revealed 19 clusters with distinguishable gene expression patterns (Fig 4b). Manual annotations were conducted to assign cell types for each cluster (Fig 4c) based on marker expression and top differentially expressed genes (DEG)(Fig 4e). We assigned the clusters as either cancer cells (C4, C9; expressing *Gpa33* and *Epcam*), NK/T-cells (C5, C15; expressing *Ptprc/CD45, CD3d, Cd3e, Cd3g* or *Nkg7*), B-cells (C7; expressing *Cd19* and *Cd79a*), endothelial cells (C13; expressing *Pecam/CD31* and *Igfbp7*), cancer-associated fibroblast (C18; expressing *Col3a1, Sparc*, and *Fstl1*), myeloid clusters (C1-4, C8, C9, C11, C12, C14, C16, C17, C18, C19; expressing *Ptprc/CD45, Itgam/CD11b, Cd68,* and *Csf1r),* and Kupffer cells (C12; expressing *Fcna, Clec4f, Vsig4*). We also confirmed our cluster assignments with an automated cell type prediction using tool SingleR(Supp Fig 2a).

Single-cell analysis of liver metastases revealed CR-Ric-LNP treatment significantly reduced the relative abundance of malignant tumor cells (↓17.5% to 1.6%), CAFs (↓3.4% to 0.4%), and resident Kupffer cells (↓3% to 1.6%) compared to control untreated lesions (Fig 4d,f).

In contrast, the myeloid, NK- and T-cells, as well as B-cells were enriched in these samples, with B-cell proportion increased from 2.6% to 7.8% and NK/T-cells doubled from 6% to 11% in CR-Ric-LNP samples (Fig 4f).

Pseudobulk analysis of DEG from CR-Ric-LNP treated samples revealed an upregulation of pro-inflammatory macrophage signatures such as *Cd40, Cd69, Iglv2,* or *Arl5c* (Fig 4g). Further, DEG analysis identified a clear upregulation of immune-related genes such *as Rsad2, Cxcl10, Iglv2, Isg15, Tnf, Ifit1/2/3, Irf7, Usp18,* and *Gbp2/4/5* (Fig 4g), indicating a robust interferon Type I/II-driven immune response and increased antigen presentation activity induced by the treatment (Fig 5e). These genes were predominantly expressed by two treatment-induced clusters: Neuts_Ifn and Macs_Ifn clusters (Suppl Fig 2c).

**Figure 5.**
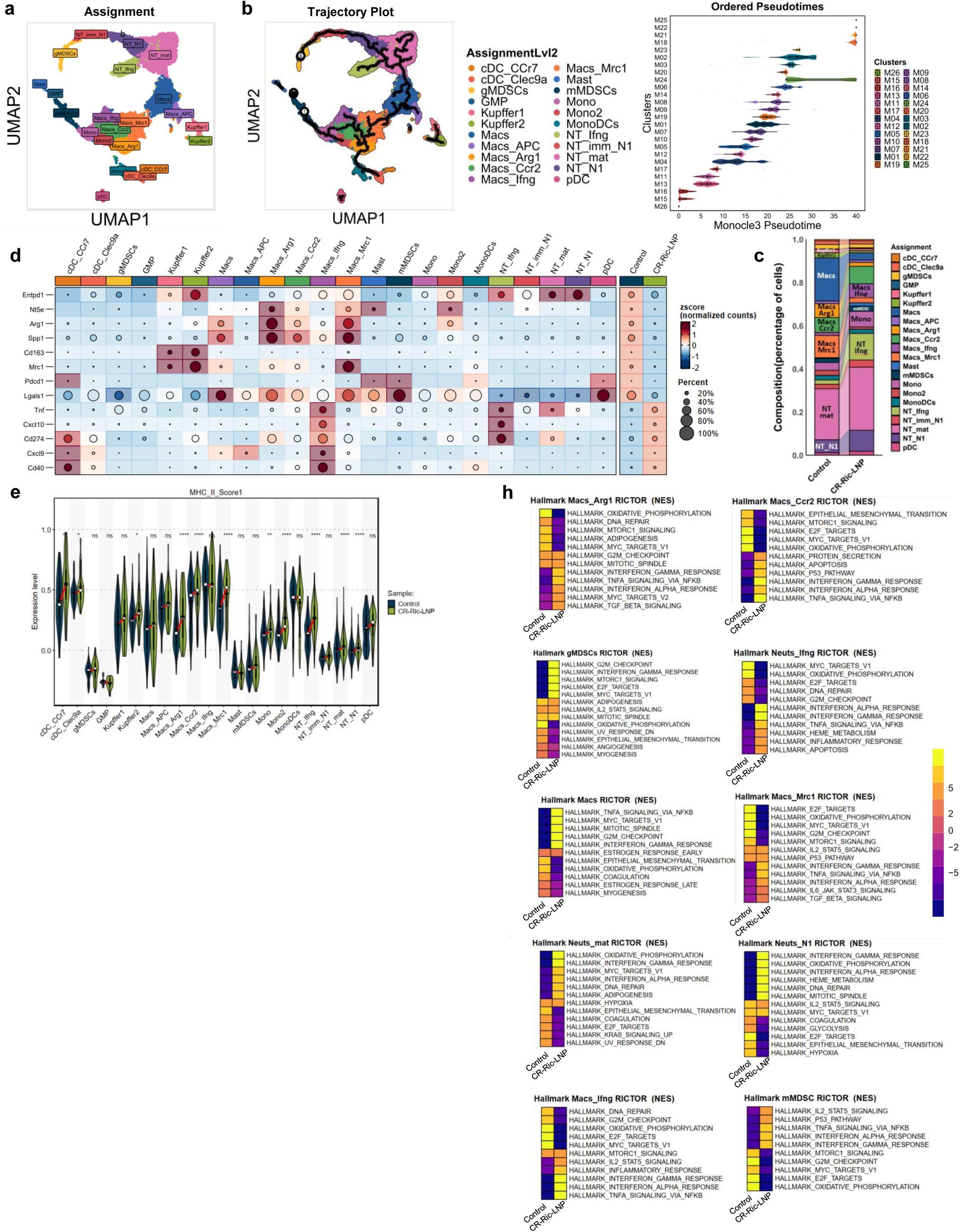
Denovo Clustering of whole Myeloid assigned clusters population revealed 26 distinct subgroups. a) UMAP projection and cell assignment of the myeloid subpopulation. b) Dimplots of pseudotime analysis of myeloid clusters revealed multiple branches for cell fates and Monocle3 pseudotime distribution trends of myeloid subsets by de novo clusters. c) Cellular composition analysis revealed a decrease of immunosuppressive myeloid cells and an increase of inflammatory subsets. d) Dot Heatmap quantification of myeloid immune checkpoint genes expressed by cell subtypes and sample. e) Violin plots show distributions of MHC-II antigen expression scores across myeloid cell clusters. Solid lines indicate interquartile ranges (IQR), with white dots representing medians. Dark green violins: untreated baseline; light green violins: with CR-Ric-LNP intervention. *p<0.05, **p<0.01, ****p<0.0001. f) GSEA Hallmark pathway analysis summary of different myeloid populations most affected by treatment.

In addition, immunosuppressive macrophage-associated genes, including *Lgals1, Ido1, CD163, Mrc1, Arg1*, *Spp1*, *Tgfa*, and the complement system genes belonging to *C1q* family exhibited significant downregulation (Fig 4g). Additionally, multiple oncogenic and stem cell-associated genes such as *Sox4, Ccn1/2/4, Igf1/2r, Wnt4,* and *Aurka/b* showed decreased expression, suggesting a broad attenuation of pro-oncogenic pathways (Fig 4g).

To evaluate the balance between proinflammatory and immunosuppressive signaling within the TME, we adapted a previously validated approach from a head and neck cancer study^76^, which used Cxcl9:Spp1 ratio as a metric for immune polarization. Given the predominance of *Cxcl10* over *Cxcl9* in our dataset, we modified this metric to evaluate the Cxcl10:Spp1 ratio. *Cxcl10*, a Th1-associated chemokine linked to anti-tumor immunity^77^, serves as a marker for immune activation and prognostic factor for response to immunotherapy in cancer^78^, including in TNBC^79^. In contrast, Spp1 (osteopontin) is associated with myeloid-driven suppression and tumor progression^80,81^. We calculated Cxcl10:Spp1 ratio across myeloid subsets and in total cells to quantify the net immunostimulatory vs suppressive signals. This analysis revealed that CR-Ric-LNP treatment significantly skewed the TME towards a pro-inflammatory state (Fig 4h, Suppl Fig 2d). Normalized Cxcl10:Spp1 ratio was elevated in total cells (Fig 4h), mirroring the predictive value previously established in head and neck cancer.

### De novo clustering of myeloid population

Next, we performed unsupervised clustering of myeloid cells, resulting in 26 clusters (Fig 5a). We conducted detailed subpopulation analysis of the myeloid subtypes based on known gene expression markers (Supp Fig 2b).

Following high resolution UMAP annotation, we delineated 5 distinct myeloid populations: neutrophil, macrophages, DC, mast, and Kupffer cells (Fig 5a). The Kupffer cells were identified and segregated from remaining myeloid cells by their characteristic expression of *Fcna, Clec4f, Cd5l* and *Vsig4* (Fig 5a, Suppl Fig 2b). Within the granulocyte subsets, reclustering revealed five transcriptionally distinct cell types: gMDSC, immature “N1” neutrophil, N1 neutrophil, mature neutrophil, and Ifn-response neutrophil cluster. For neutrophil assignments, we adopted the maturation-based nomenclature as previously described^82^. Thus, the designation of “N1” refers strictly to the developmental stage of such cells, rather than their inflammatory or activation state. In parallel, the monocytic lineage also had multiple subsets with varying maturation states. The lineage originating from the common GMP progenitor shared with neutrophil population, progresses through classical monocytes, to mature macrophages, with 5 major subtypes, annotated as Macs_Ccr2, Macs_Arg1, Macs_Mrc1, Macs_Ifn, and Macs.

Our analysis revealed a marked decrease in immunosuppressive macrophage subtypes, particularly within the Macs_Mrc1, Macs_Arg1, and general Macs clusters in CR-Ric-LNP treated metastases. Concurrently, there was notable downregulation of immunosuppressive macrophage markers, including *Arg1, Nt5e, Entpd1, Cd163, Pdgfa, Mrc1, Apoe, Ccl6* and *C1qa/b/c*.

We also identified treatment-induced Ifn-I/II*-*primed subpopulations within both neutrophils (Neuts_Ifn) and macrophages (Macs_Ifn). These clusters exhibited activation of *Fcgr1* (CD64), a hallmark of Ifn-primed phagocytes, enrichment for *type I interferon* (*Ifnb1*, *Irf7*) and *type II interferon* (*Ifngr1*) signaling pathways, increased expression of *Isg15, Ifit1*, and *Rsad2* (Fig. 5a, Supp Fig 2b,c). The pseudotime trajectory from these clusters revealed that granulocytic monocyte progenitor (GMP) serving as the origin, before diverging into two distinct branches: one differentiating into gMDSC and neutrophil, and the other one progressing towards mMDSC and monocytic cell subsets before ultimately transitioning into mature macrophages and DCs (Fig 5b). CR-Ric-LNP treatment decreased Macs, Macs_Arg1, and Macs_Mrc1 from 20%, 6.4%, and 10.3% of myeloid cells in control samples to 3.3%, 1.8%, and 2.3% in treated cohorts, respectively (Fig 5c). In contrast, both Macs_Ccr2 (from 6.95 to 8.1%) and Macs_Ifn populations (from 1.4% to 6.5%) were increased in CR-Ric-LNP treated liver metastases.

Neutrophils were generally increased: Neuts_Imm_N1, Neuts_N1, and Neuts_mat were moderately increased from 6%, 2%, 23.6% in control to 9.7%, 3.3%, and 29.2% in CR-Ric-LNP treated samples, respectively (Fig 5c). The *Ifn*-response neutrophil cluster increased 6.2-fold, rising from 2.0% in controls to 12.3% in treated samples, marking the most pronounced compositional shift in the myeloid compartment (Fig 5c). Single-cell compositional analysis revealed resilience of dendritic cell (DC) populations to treatment. Conventional DC subsets (cDC1, cDC2) remained unchanged, with only plasmacytoid DCs (pDCs) showed an increase from 1.2% in control to 1.8% in treated samples (Fig. 5c).

Analysis of immune checkpoint gene expression confirmed distinct phenotypic and functional profiles across cell types. The Neuts_Ifn and Macs_Ifn subsets exhibited elevated expression of pro-inflammatory mediators (*Cxcl10*, *Tnf*) and the immune checkpoint gene *Cd274* (PD-L1), with Macs_Ifn uniquely displaying robust *Cxcl9* and *Cd40* activation, a co-stimulatory signal critical for antigen presentation, which only shared with cDC_Ccr7 population (Fig. 5d, Supp Fig 2c). Both Ifn-primed populations are also the major contributor for *Cxcl10* expression (Supp Fig 2c). Conversely, immunosuppressive macrophage subsets Macs_Mrc1 and Macs_Arg1 were enriched in checkpoint-associated molecules such as *Vsir* (VISTA), *Nt5e* (CD73), and *Spp1* (osteopontin) (Fig 5d), which are linked to immune evasion. To assess antigen-presenting capacity, we evaluated MHC class II (MHC-II) gene expression across myeloid clusters using a scoring metric based on canonical MHC-II genes (Table 2). Nearly all myeloid clusters exhibited elevated MHC-II scores including DCs, inflammatory macrophages, and monocytes (Fig 5e).

**Table 1.** Antibodies for IMC staining.

|  | Phenotyping marker |
| --- | --- |
| Cytotoxic lymphocytes | CD8 |
| T helper | CD4 |
| Macrophages | F4/80 |
|  | CD86 |
| M2 macrophage | CD163 |
| NK Cells | NK1.1 |
| Dendritic cells | CD11b/CD11c |
| Cancer cells | Ki67/PanCK |
| MDSC (granulocytic) | Ly6G |
| Endothelial cells | Endomucin |
| B-cell | B220 |
| Treg | Foxp3 |

| Functional markers |
| --- |
| PD-1 |
| PD-L1 |
| GATA-3 |
| CD44 |
| pSTAT1 |
| pSTAT3 |
| pERK |
| p-Akt |
| p-p38 |

**Table 2.** Gene sets used for composite scoring. Inflammation scores reflect pro-inflammatory states; MHC-I/II scores represent antigen presentation capacity.

| Inflammatory | MHC_I | MHC_II |
| --- | --- | --- |
| *Ifng* | *H2-K1* | *H2-Aa* |
| *Tnf* | *H2-D1* | *H2-Ab1* |
| *Il1b* | *B2m* | *H2-Eb1* |
| *Il6* | *Tap1* | *Cd74* |
| *Cxcl9* | *Tap2* | *Ciita* |
| *Cxcl10* | *Psmb8* | *H2-Oa* |
| *Nos2* | *Psmb9* | *H2-DMa* |
| *Stat1* | *Tapbp* | *H2-DMb1* |
| *Irf1* | *Stat1* | *Irf8* |
| *Ccl5* | *Irf1* | *Stat1* |
| *Ccl2* | *Calr* | *Cd86* |
| *Nfkb1* | *Canx* | *Cd40* |
| *Ccl3* | *Erp57* |  |
| *Ccl4* |  |  |
| *Gbp2* |  |  |
| *Il12b* |  |  |
| *Batf2* |  |  |
| *Irf7* |  |  |
| *Gbp5* |  |  |
| *Cd80* |  |  |
| *Cd86* |  |  |

GSEA Hallmark pathway analysis revealed consistent positive enrichment of the Interferon_Alpha/Beta_Response and TNFα_Signaling_via_NFκB pathways across all myeloid clusters, irrespective of their immunosuppressive states, suggesting broad type I/II IFN and inflammatory signaling activation in response to therapy (Fig. 5f). Notably, Macs_Ifn and Neuts_Ifng exhibited negative enrichment for Oxidative_Phosphorylation (OXPHOS) post CR-Ric-LNP therapy (Fig 5f), indicating a metabolic shift away from mitochondrial respiration and a shift towards glycolysis, associated with pro-inflammatory (M1-like) activation^83^, correlated to favorable metabolic rewiring toward immune activation and tumor cell killing^84^.

In contrast, mature neutrophils showed significant OXPHOS upregulation post-treatment, highlighting a divergent metabolic response between myeloid subsets. This suggests that CR-Ric-LNP therapy differentially reprograms cellular metabolism, potentially influencing their functional states in the TME.

### Effects on T-cells, B-cells and CAFs

To assess treatment-induced shifts in T-cell composition, we performed *de novo* reclustering of T cells. Cell identities were annotated using canonical gene markers (Fig. 6a) and visualized via UMAP projection (Fig. 6b). Striking changes in subset distribution emerged: exhausted CD8⁺ T cells and regulatory T cells (Tregs) were markedly reduced, whereas NK cells, Ifn-primed CD8⁺ T cells (*CD8_Ifn*), and naïve CD8⁺/CD4⁺ T cells expanded significantly (Fig. 6c).

**Figure 6.**
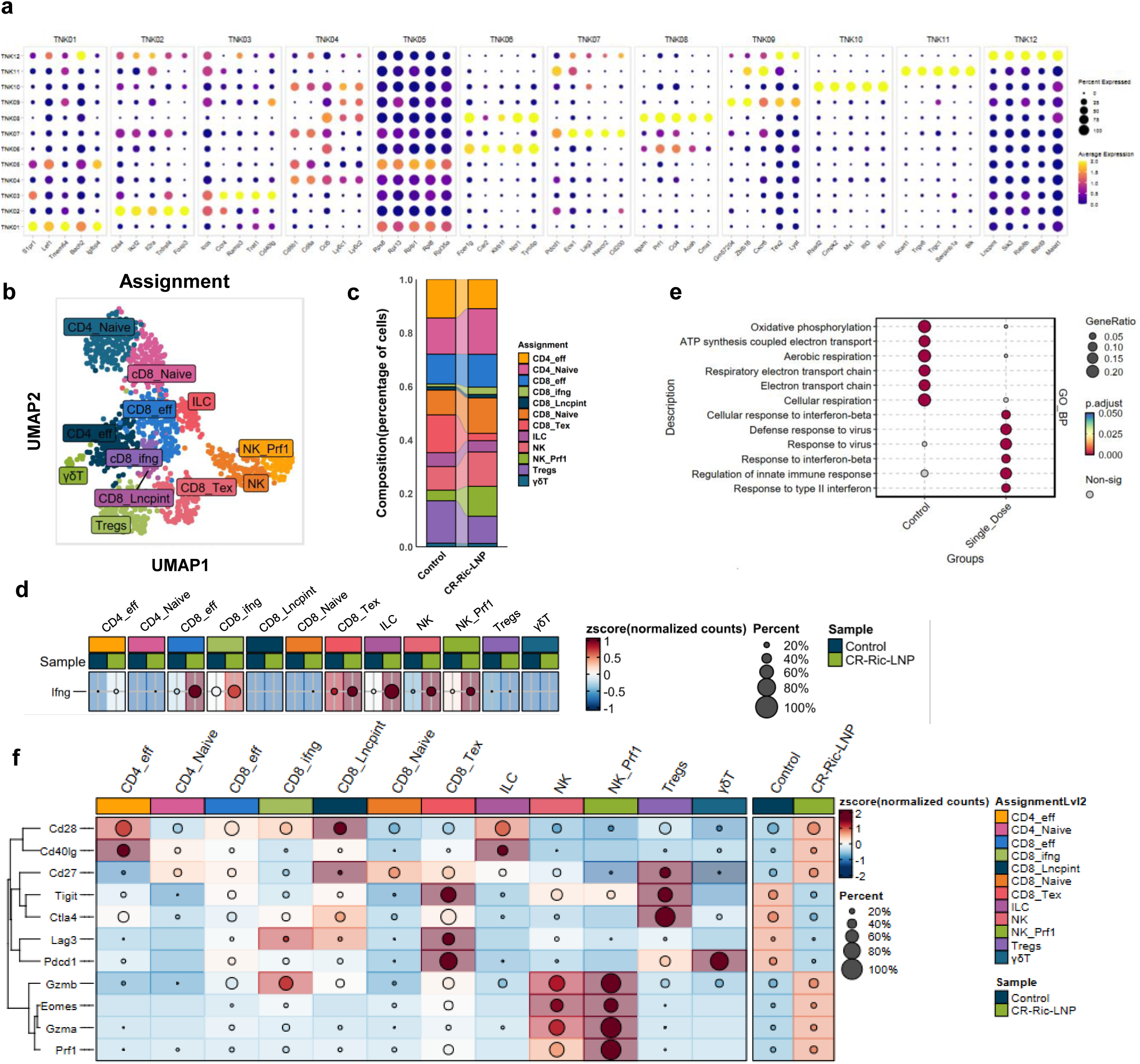
Denovo Clustering of T cells and NK cells assigned clusters population revealed 12 distinct subgroups. a) Dot plot for the top differential genes of T-cell associated clusters for subpopulation assignments. Dot size indicates the fraction of expressing cells, colored based on normalized expression levels. b) UMAP visualization of T-cells with 12 distinct clusters, c) Quantification of T-cell subpopulation within the cluster showing significant shift of T-cell subpopulations after CR-Ric-LNP treatment. d) Dot Heatmaps depicting the shift in *Ifng* (IFNγ) expression levels among T cell subpopulations following treatment with CR-Ric-LNP. Each dot represents a T cell cluster, with color intensity (gradient: blue → red) indicating the average *Ifng* expression level and dot size reflecting the percentage of cells expressing *Ifng* within the cluster. e) Gene Ontology Biological Process (GO_BP) analysis of differentially expressed pathways in T-cell clusters following treatment. Dot plots display the top **upregulated** (red) **GO_BP** in T cell clusters across samples. f) Heatmaps indicating normalized expressions of select immunomodulatory and immune checkpoint genes in T-cell clusters and by sample.

NK cell proportion within T-cell cluster increased 1.5-fold (8.8% to 12.9%), and Prf1⁺ cytotoxic NK cells surged nearly 3-fold (4% to 11.2%) in treated cohorts. Conversely, exhausted CD8⁺ T cells declined >5-fold (14.2% to 2.7%), and Tregs decreased 1.5-fold (15.8% to 10.1%) post-treatment (Fig. 6c). Naïve CD4+ T-cells increased from 13.5% to 17.1%, naïve CD8+ T-cells from 9.3% to 13.3% in treated samples. These dynamics suggest treatment-driven enhancement of cytotoxic and effector functions, coupled with attenuation of exhaustion and immunosuppression. Among these T-cell populations, CD8_Ifng population, followed by innate lymphoid cells (ILC), exhausted CD8 T-cells, effector T-cells, and NK cells are the major contributor of *Ifng* expression (Fig 6d). CR-Ric-LNP treatment significantly enhanced *Ifng* expression across these populations. Gene Ontology Biological Processes (GO_BP) analysis on T-cells revealed significant alterations in both immune and metabolic pathways, such as response to interferon gamma and beta, along with increased regulation of innate immune response and response to virus (Fig 6e). Cellular respiration and ATP-synthesis coupled electron transport as well as oxidative phosphorylation processes were reduced in T-cells after treatment. Transcriptional analysis revealed significant alterations in checkpoint protein expression across T cell subsets. We observed a significant downregulation of immune checkpoint molecules including Pdcd1, Tigit, Ctla4, and Lag3 across the general T-cell population (Fig. 6f), suggesting a reduction in T-cell exhaustion and potential restoration of anti-tumor immune activity after treatment. *Pdcd1* was substantially reduced in exhausted, regulatory, and γδ T-cells, while *Tigit* showed moderate downregulation in exhausted, regulatory T-cells, and NK cells (Fig 6f). Notably, *Lag3* was moderately elevated in exhausted T-cells, while *Ctla4* expression remained unchanged in most T-cell populations, but decreased in ILC, NK cells, and in γδ T-cells (Fig 6f).

The increased chemokines for TIL infiltration was further accompanied by the increased expression of cytotoxic markers (*Gzma, Gzmb, Prf1)* and T-cell activation genes (*Cd69, Cd40, Cd3e/g, Lck, Lat, Ncr*), suggesting robust cytotoxic lymphocyte activation and tumor cell killing capacity. Expression of activated macrophage marker such as *Cd40*, along with other genes such as *Cd83* which indicates DC maturations, *Ebf1*, a B-cell expansion marker^85,86^, and *Mzb*, a factor implicated in B-cell and pDC polarization and antibody secretion^87^ suggested an increased antigen presentation activity and potential formation of immune synaptic center^88^.

Additionally, genes related to ECM stiffening, fibrosis, and metastasis, which can promote collagen deposition, crosslinking, and matrix degradation were downregulated upon treatment. These genes include *Fn1* (Fibronectin 1), *Col1a1, Col3a1, Col5a1, Col16a1, Col18a1, Sparc, Loxl3, Timp1*, and *Mmp14*. This, along with significant reduction of CAFs proportion within TME after treatment indicates reduced stromal support and impaired metastasis and angiogenesis^89,90^.

Inflammation score was increased particularly in myeloid cell population (Fig 7a), and MHC-I scores were increased in multiple populations including myeloid, T/NK cells, and B-cells (Fig 7b). CellChat analysis of tumor-immune interactions revealed profound changes in inferred cellular communication networks following CR-Ric-LNP treatment in liver metastasis samples. In the untreated samples, the predominant interactions involved tumor and immunosuppressive Macs_Arg, Macs_Mrc1, and Macs_Ccr2 (Fig 7c). CR-Ric-LNP treatment showed disruption of this immunosuppressive network, with weakening immunosuppressive Macs-tumor cell crosstalk. Signals from exhausted CD8 T-cells and CAF were also diminished, suggesting diminishing angiogenesis and fibrotic processes.

**Figure 7.**
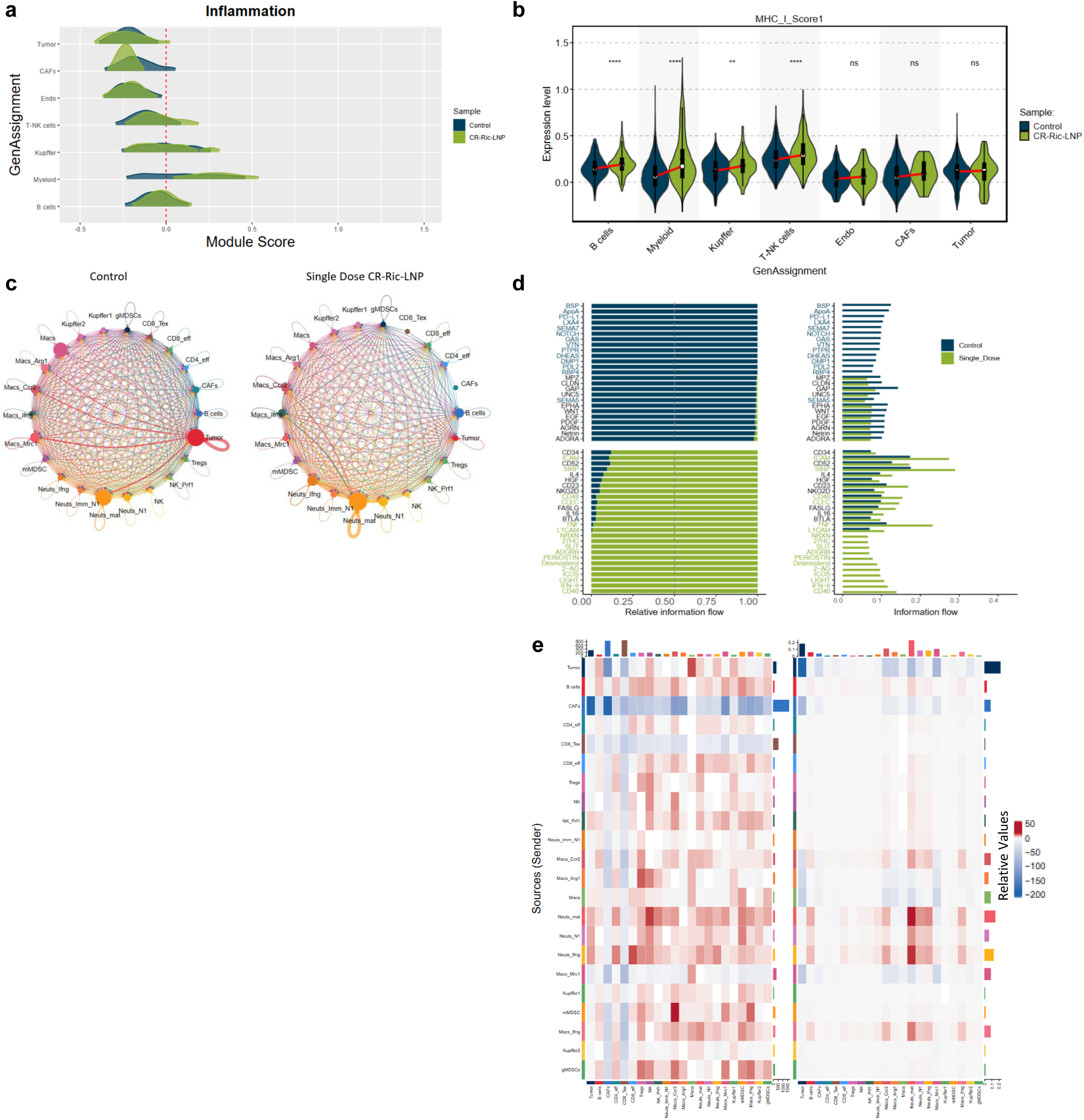
Inferred cell-cell interaction from Cellchat analysis of scRNAseq data revealed global signaling changes between Control and Rictor KD. a) Density distributions of inflammation scores and MHC-I scores across cell clusters split by the respective sample distribution. Scores are derived from the mean expression of pro-inflammatory and MHC-I genes (Table 2). c) Circular plot of total weighted interaction, a measure of significance based on ligand, mediator, and receptor expression, are summarized across cell types. d) Top pathways are displayed in an information flow plot to highlight the shift in cellular interactions pathways between control and CR-Ric-LNP treated samples. e) Top 25 enriched pathways in control and in Rictor KD samples in regards of relative and absolute information interaction strength. e) Differential heatmap for number of interactions between cell clusters in the TME. The vertical axis represents the signal sender, and the horizontal axis represents the signal receiver. The blue column indicates decreased communication in the Rictor KD samples compared to that in the control, while red indicates increased communication.

Further investigation identified key genes activated from the cell-cell interactions, where the IFN-II signaling was dominated by exhausted CD8 T-cells in control tumor, the Ifng signaling originated predominantly from NK and effector CD8 T-cell populations in treated samples (Suppl fig 3d). Surprisingly, the top enhanced pathways for cellular interactions include those linked to B-cell (*Btla, 2-AG*) (Fig. 7d). Outgoing and incoming signaling patterns demonstrated a clear reduction in tumor-derived signals, including oncogenes and genes mediating intercellular crosstalk (Fig. 7e). These findings also suggest that CR-Ric-LNP treatment disrupts immunosuppressive tumor-immune interactions while promoting a more active immune response. Evaluation of immunomodulatory pathways revealed activation of MHC-II, TNF, and IFN-II pathways (Suppl Fig 3a,b,c).

CR-Ric-LNP induced a broad transcriptional downregulation in tumor cells, as evidenced by negative enrichment across nearly all GSEA Hallmark pathways (Suppl Fig 4a). Only modest enrichment was observed in Hallmark_Kras_Signaling_DN, suggesting partial downregulation of Kras-driven oncogenic signaling. Other non-malignant populations significantly affected by the treatment are endothelial cells and CAFs (Suppl Fig 4b,c). Both populations exhibit parallel negative enrichment for most GSEA Hallmark pathways, suggesting reduced stromal and vascular support for tumor growth. Slight enrichment in bile acid metabolism pathway for both populations may reflect to lipid-rich environment, which is relevant in the liver metastasis context.

In contrast, B-cell displayed distinct enrichment dynamic. Most of GSEA Hallmark pathways were positively enriched, with only several negative tumor-related pathways, such as IL_Jak_Stat3, angiogenesis, EMT/myogenesis, or coagulation, hypoxia, and inflammatory response (Suppl Fig 4d).

### Validation of scRNAseq findings via IMC and immunofluorescence confirms enhanced anti-tumor immunity

To validate our scRNAseq results at the protein level, we performed imaging mass cytometry (IMC) and immunofluorescence (IF) analysis of metastatic liver lesions (0.25–1 mm³, n=6/group). IMC was conducted using a 21-plex antibodies (Table 1). Data from IMC analysis showed increased immune cell infiltration within tumor lesions. CD8+ T cell infiltration increased, including PD-1+ CD8+ T cells in close proximity to macrophages within metastatic lesions (Fig 8a, b, c). Elevated CD8+ population was also observed from the IF-stained samples after treatment with CR-Ric-LNP (Fig 8c). Neighborhood analysis showed that CD8+ cells in the tumor vicinity increased from 2.7% and 1.9% the control samples within 30μm and 100 μm around tumor, respectively, to 8.2% and 8.7% in CR-Ric-LNP treated samples (Fig 8b). Strikingly, CR-Ric-LNP treatment led to a nearly 4-fold reduction in Ki-67+ proliferating tumor cells compared to scrambled CR-LNP controls after a single dose (Fig. 8d).

**Figure 8.**
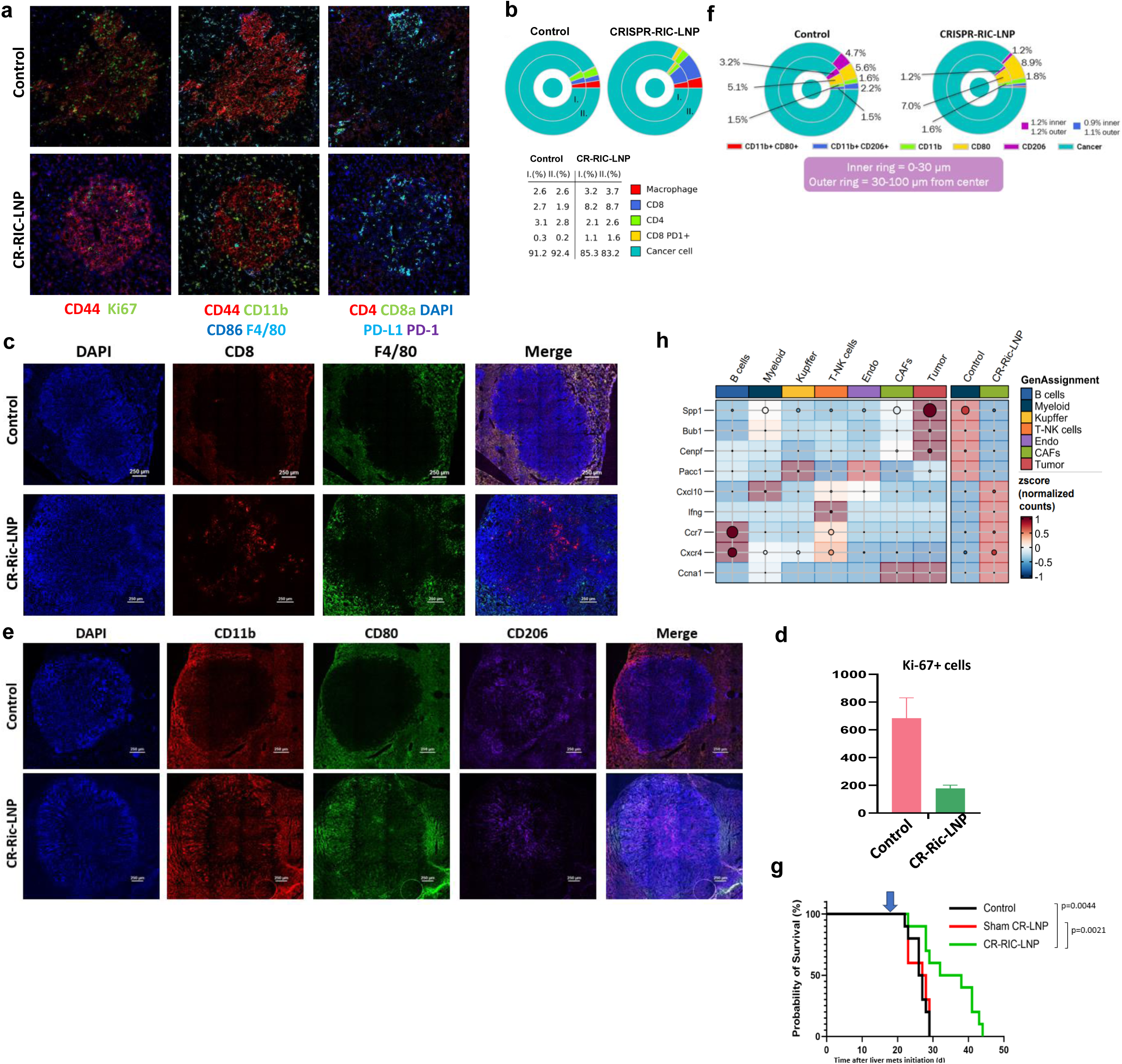
In vivo evaluation of CR-Ric-LNP efficacy via Imaging Flow Cytometry (IMC), cellular staining, and survival study. a) IMC analysis of liver metastases from 4T1 murine model stained with antibodies for cellular functions and T-cell exhaustion markers b) CR-Ric-LNPs treatment reduced Ki-67 expression in metastatic lesions of 4T1 murine livermetastasis model as detected via IMC. c) Representative immunofluorescence images of liver metastases stained with CD11b, CD80, CD206 antibodies and counterstained with DAPI. d) Composition analysis from liver sections stained via immunofluorescence markers for macrophage subtypes CD11b, CD80 and CD206. e) Representative immunofluorescence images of liver metastases stained with CD8, and F4/80 antibodies, and counterstained with DAPI. f) Composition analysis from liver sections stained for pan macrophage (F4/80), T-cells (CD4, CD8), exhaustion (PD1), and cancer (PanCK) markers. Blue, yellow and red doughnut slices denote macrophages, CD8+ T cells, and CD8+ PD1+ T cells, respectively. g) Kaplan-Meier curve of survival study after one-time treatment with CR-RIC-LNP (d18, blue arrow) in 4T1 BCLM model. h) Dotplot for published TNBC prognostic factors gene expression, cellular clusters expressing the genes, and changes in TNBC-LM model after CR-Ric-LNP treatment.

Quantitative analysis of IF staining of whole liver sections using Qupath (Suppl Fig 5a) demonstrated a shift in macrophage accumulation. We observed an increase of CD11b+ and CD80+ signal in the liver metastases treated with CR-Ric-LNP (Fig 8e). Quantification of the signal indicating that CD80-expressing pro-inflammatory macrophages increased from 5.1% to 7% in direct contact with tumor cells and from 5.6% to 8.9% in the surrounding 100µm TME (Fig 8f). Conversely, CD206+ immunosuppressive macrophages significantly decreased from 3.2% to 1.2% within 30µm of tumor cells and from 4.7% to 1.2% in the broader tumor-adjacent regions (Fig. 8f).

The observed immunological remodeling translated into a robust survival benefit in the aggressive 4T1 liver metastasis model. A single dose of CR-Ric-LNP extended median survival from 26 days (Sham CR-LNP control) to 38 days (CR-Ric-LNP group), representing a 46% increase in lifespan (p=0.0021, Fig. 8g).

We further evaluated the changes of gene expression of various genes previously reported as prognostic factors in TNBC^79,91^. The dot plot illustrates the impact of a macrophage-modulating treatment in breast cancer, revealing significant downregulation of *Spp1, Bub1, Cenpf*, and *Pacc1* alongside upregulation of *Cxcl10, Ifng, Ccr7, Cxcr4*, and *Ccna1* (Fig 8g). The reduced expression of *Spp1* (osteopontin) and mitotic regulators (*Bub1, Cenpf*) suggests diminished tumor aggressiveness and genomic instability, while *Pacc1* downregulation may disrupt tumor acidosis, a key immune evasion mechanism. Conversely, elevated *Cxcl10* and *Ifng* indicate robust anti-tumor immunity, recruiting cytotoxic T cells and polarizing macrophages toward an M1 phenotype. *Ccr7* upregulation further supports adaptive immune priming, though *Cxcr4* role remains context-dependent—potentially enhancing immune trafficking but requiring caution due to its metastatic links. The increase in *Ccna1*, a cell cycle marker, may reflect proliferative immune activation. Collectively, these shifts align with improved prognostic signatures in breast cancer, including reduced metastasis (*Spp1↓*), enhanced immune infiltration (Cxcl10↑/Ifng↑), and better survival outcomes (Bub1↓/Ccr7↑) (Fig 8g), highlighting the therapy’s dual role in suppressing tumorigenic pathways while fostering an immunostimulatory microenvironment.

Histopathological assessment revealed no evidence of hepatotoxicity in Balb/c mice following either single or repeated CR-Ric-LNP administration. Liver architecture remained intact, with preserved lobular organization and absence of necrosis or fibrosis and no signs of degeneration in hepatocytes morphology (Suppl Fig 5b).

## Discussion

We developed a CRISPR-LNP system in this study, which offers a novel approach to modulating macrophage phenotypes for therapeutic options. Cpf1/Cas 12a was selected as CRISPR nuclease over Cas9 for present study for these reasons: 1) Cpf1 leaves short overhangs as sticky ends, allowing more efficient and accurate DNA editing; 2) protospacer adjacent motif (PAM) for Cpf1 is more abundant in the genome compared to the G-rich PAM targeted by Cas9; and 3) Cpf1 requires only a single RNA (∼42 nt long) as compared to two small RNAs (∼100 nt crRNA/tracrRNA) for Cas9, enabling easier loading into LNP. We have successfully produced multiple batches of LNP with minimal batch variation, demonstrating robust generation and readiness for Good Manufacturing Practice (GMP) translation for future preclinical studies.

Upon uptake into the cells, the CR-LNPs are processed rapidly. The internalized particles containing ionizable lipids becomes protonated upon entering low pH condition within endosomes, allowing them to interact with endosomal membranes and facilitate the release of encapsulated CRISPR content into the cytoplasm^92^. Following this release, the nuclear localization sequence (NLS) within the Cas12a nuclease directs its relocation to the nucleus, where CRISPR activity can be detected by the presence of double strand breaks, some occurring within just 5 minutes after administration (Fig 1c), consistent with our previous reports with similar Cas9-encapsulating CR-LNP^93,94^.

In our earlier work, we showed that the CR-Ric-LNP treatment effectively downregulates *Rictor* expression in BMDMs, with Western blot analysis confirming a marked reduction in *Rictor* protein levels^62^. Current findings also demonstrate that although CR-Ric-LNP effectively induced genetic reprogramming in macrophages, treatment did not cause significant alter naïve, unstimulated macrophages (Fig 2c). This effect confirms the non-immunogenic nature of the CR-LNP platform itself, underscoring the favorable safety profile for therapeutic applications.

Biodistribution data revealed that upon injection, the majority of the LNPs were taken up by circulating CD11b+ myeloid cells, including neutrophils and monocytes, which were subsequently recruited into the metastatic lesions in the liver (Fig 3). This is a significant achievement, as penetrating liver metastases is notoriously difficult due to the typical hypovascularized nature of tumors in breast cancer liver metastases^95^. Our previous studies indicated that the capillaries within the metastatic lesions are underdeveloped, and the internal pressure of the lesions further hinders the penetration of macromolecules into the metastatic sites^75^. A portion of the ionizable LNPs also inherently accumulated in the liver, as is typical for ionizable LNP and has been previously reported^96^. Particles were cleared from uninvolved liver within 4 hours after treatment, remaining only in the tumor lesions.

Our scRNAseq data indicates that the penetration of reprogrammed myeloid cells into liver metastases was able to remodel the TME 1 day after administration of CR-Ric-LNP. Treatment induced significant shifts in cellular composition, aligning with the expected anti-tumor immune response and tumor cell killing (Fig 4f). In the TME, increased immunosuppressive M2 polarization is the leading cause of T-cell exhaustion^97^, negatively impacting the immune system ability for tumor clearance. M2 macrophages can contribute to immune exhaustion by reducing available amino acids such as L-arginine and tryptophan thereby inhibiting CD8+ cytotoxicity^98^.

Our data revealed a significant reduction in both the Macs_Arg1 population and global *Arg1* expression, a key enzyme that depletes L-arginine and generates immunosuppressive ornithine in the TME. This reduction was accompanied by downregulation of *Nt5e/Cd73* and *Entpd1/Cd39,* critical ectoenzymes responsible for adenosine production, another major immunosuppressive mediator^99^. The coordinated suppression of *Arg1*-mediated L-arginine metabolism and the CD73/CD39-adenosine pathways, both well-established drivers of T cell dysfunction and myeloid suppression, led to increased inflammatory characterization in myeloid population (Fig 5e), which strongly indicates a breakdown of immunosuppressive networks in the TME.

De novo clustering analysis of T-cells also revealed increases of *Ifng* expression in T-cells, particularly in activated T-cell clusters (Fig 6d). Furthermore, we also observed the switch of Ifn-II signaling from exhausted T-cells in control tumor, associated with dysfunctional signaling to effector T-cells and NK cells in treated samples (Suppl fig 3c), enhancing *Cd274* (PD-L1) expression in T-cells and Ifn-primed myeloid clusters (Fig 5d, 6f) for immune checkpoint priming. Previously, high IFN-response signature has been correlated with response to immunotherapy, such as anti-PD-1 in melanoma and lung cancer^100,101^, suggesting similar potential in our platform. Genes related to chemokine and cytokine production such as *Cxcl10, Ccl3/4/5, Tnf, Saa3* were highly upregulated after treatment, supporting chemotactic recruitment of TILs. High expression of *CXCL10* has been reported as prognostic factor of increased survival in TNBC^79^, advanced serous ovarian cancer^102^ and metastatic melanoma^78^, as well as response to neoadjuvant chemoradiotherapy in rectal cancer patients^103^.

In contrast, expression of *Spp1*, a major protein correlated with tumor progression, metastasis and resistance to therapy^76,81,104^, was significantly reduced upon CR-Ric-LNP treatment *in vivo*. Pro-tumorigenic genes such as *Lgals1* (Galectin-1), *Thbs1* (Thrombospondin 1), *Serpine1* (PAI-1), *Wnt7b*, and *Bmp7* were downregulated, suggesting a reduction in tumor cell survival, angiogenesis, and immune evasion.

Upregulation of MHC-II in myeloid population enhances the presentation of antigens to CD4+ T cells, facilitating improved T-cell activation and proliferation, as also shown in reduction of expression of immune checkpoints crucial for regulating immune activity and preventing autoimmunity such as *Pdcd1/*PD-1, *Ctla4, Tigit* and *Lag3* in T-cells. Additionally, treatment with CR-Ric-LNP also increased the reservoir of naive CD8+ T-cells. This represents a “fresh army” which can be primed for activation with additional interventions. Global T-cell analysis also revealed elevated immune response, suggesting a crosstalk with myeloid cells and induction of pro-inflammatory T-cell phenotype, consistent with our observed TME remodeling towards more “hot” state. Thus, the reduction of cellular respiration and ATP-synthesis coupled electron transport from GO_BP analysis of T-cells implicated a shift away from mitochondrial metabolism, typical of highly activated or effector T-cells reliant on glycolysis.

We found that despite a strong inflammatory response, there was some immunosuppressive resistance that limit the inflammation, such as Macs_Arg1 population. This potentially derived from biological feedback mechanisms to protect from excessive inflammation and tissue damage. Additionally, the tolerogenic environment in the liver may also resist full immune activation in effort to protect the liver from autoimmune hepatitis or fibrosis.

Among the population implicated in the TME, our data also suggests enhanced proliferation and changes in B-cells properties along with reprogramming and diminishing numbers of CAFs populations, highlighting the need for future in-depth studies. B-cells have the potential for antibody production and serve as a source of plasma cells or tertiary lymphoid structure that correlates with better prognosis for patients. In contrast, treatment led to a significant shift in CAFs functional state, implicating changes in metabolism and therapy response, indicating reduced CAF-mediated stromal support for cancer cell survival and metastasis.

This transcriptomic data was consistent with the data observed from protein level from IMC and IF analysis, showing increased inflammatory and cytotoxic myeloid and T-cell expansion, leading to reduced Ki-67+ cells in CR-Ric-LNP treated liver metastases (Fig 8c). We also observed a trend in increasing CD8+ T-cell and macrophages population, including the increased PD-L1 expression in the treated samples.

Overall, our findings suggest a secondary immunostimulatory mechanism triggered by the depletion of immunosuppressive macrophages. As Macs_Mrc1 and Macs_Arg1 populations declined, we observed a concomitant increase in IFNγ secretion from T cells, which in turn promoted the expansion of Ifn-primed myeloid clusters (Neuts_Ifn and Macs_Ifn). This self-reinforcing feedback loop, where reduced immunosuppression enhances T cell-derived IFNγ, further amplifying the recruitment and activation of interferon-sensitive immune cells, which highlights a key mechanism by which the therapy sustains an anti-tumor immune response.

The dynamic shift from an immunosuppressive to an IFNγ-driven inflammatory microenvironment underscores the multi-layered efficacy of our CR-Ric-LNP treatment, which enhanced both innate and adaptive immune responses to orchestrate a broader immune-activating cascade to obtain a significant survival benefit. By modulating checkpoint molecules and increasing antigen presentation, CR-Ric-LNP holds promise for improving therapeutic outcomes in treating infections and tumors, potentially leading to more effective immune-mediated eradication of disease cells.

Further, we observed statistically significant improvement of survival (p=0.0021) which suggests that even transient reprogramming of the TME can produce clinically useful outcomes in advanced metastatic disease. Analysis of gene expression previously reported genes as potential prognosis factors in TNBC indicated a strong positive trend towards increased survival (Fig 8f). Increased *Cxcl10, Cxcr4*^79^, and *Ccr7*, as well as reduction of *Bub1* and *Pacc1*^91^ observed after CR-Ric-LNP treatment were correlated with increased rates of disease-free survival^91^. *Spp1* gene expression was shown to be strongly reduced from CR-Ric-LNP treatment compared to the untreated control. This gene is associated with immunosuppressive macrophage infiltration and poor prognosis in TNBC^104^.

The magnitude of survival extension is particularly notable given that the treatment was administered as monotherapy in late-stage metastasis in the 4T1 model, which is highly aggressive and resistant to most interventions. Additionally, benefits were achieved with just a single therapeutic dose. These results strongly support further development of CR-Ric-LNP, particularly when considering its favorable safety profile observed in previous toxicity assessments.

## Future directions

We are presenting promising data that demonstrates efficient TME reprogramming and an overall survival benefit with our CR-Ric-LNP therapy for breast cancer liver metastasis.

Although the present study provides a strong premise for the potential of CR-Ric-LNP therapy, longitudinal studies are necessary to assess changes in the TME over extended periods. While therapy may increase survival benefits in late-stage liver metastasis models, the animals eventually succumb to the disease, indicating that repeated doses may be required to maintain a sufficient inflammatory milieu in the TME to support ongoing anti-tumorigenic activities. Our findings suggest that complementary strategies targeting myeloid cell functions, particularly through enhancement of phagocytic activity via SIRPα-CD47 axis blockade could further augment tumor cell elimination and therapeutic efficacy. Previously, CD47 inhibitors (e.g., magrolimab) activity to enhance macrophage phagocytosis of tumor cells relies on the activation of T-cells and the presence of T-cell-derived IFN-γ^105,106^, aligning with our observed T-cell activation.

The ability of CR-Ric-LNP to remodel the myeloid compartment and reinvigorate T cell function presents an attractive opportunity for combination with existing immune checkpoint inhibitors (ICIs). Not only that this approach aligns with but may also overcome limitations in prior clinical efforts to target innate immunity.

In clinical trials NCT02713529 and NCT02880371, anti-colony-stimulating factor 1 receptor (anti-CSF1R) monoclonal antibody and inhibitor were tested in combination with pembrolizumab. Both studies showed acceptable safety profiles of anti Csf1r in multiple solid cancers, albeit with limited anti-tumor activity^107,108^ due to compensatory activation of regulatory T cells^109^ and increased recruitment of other suppressive myeloid populations^110^. In another study, a CD40 agonist (sotigalimab) has been tested in combination with nivolumab in a Phase II trial, showing a favorable safety profile with a consistent therapeutic response^111^. Combination of CD40 agonist, CDX-1140, and pembrolizumab has been evaluated in multiple solid cancers, with acceptable safety profile and evident clinical benefit in patients with squamous cell carcinoma of head and neck, warranting further studies in trial NCT03329950^112^.

Notably, our data demonstrated substantial TME remodeling towards an immune-responsive profile characterized by increased baseline intra-tumoral infiltration of naïve T-cells and reduced proportion of exhausted and regulatory T-cells. Altogether, this suggests successful reprogramming of the TME from a “cold” tumor to a “hot” environment, which is more responsive to immunotherapy, indicating that this therapy may synergize other immunotherapies like anti-PD-1, PD-L1, CTLA-4, or Lag3 therapies.

However, further investigation is required to address several critical questions. Future studies should assess the exact population directly affected by the treatment, how long the pro-inflammatory shift persists post-treatment and whether sustained immune activation can be achieved. Systematic evaluation of dosage and frequency will be essential to determine the regimen required for durable anti-tumor immunity. Such studies will be crucial for translating CR-Ric-LNP into a clinically viable strategy, either as a monotherapy or in combination with existing immunotherapies.

## Materials and Methods

### Formulation and Assembly of ionizable Lipid Nanoparticles (iLNP)

The CRISPR-LNPs were generated through the hydration-extrusion method as previously described^62^. In brief, Phospholipids and cholesterol materials were dissolved in ethanol, with fluorescent phospholipid Lissamine rhodamine B DPHE, triethyl-ammonium salt (Invitrogen, Carlsad, CA) added for biodistribution studies.

A thin film of LNP material was deposited when the ethanol evaporated after 30 minutes on a rotary evaporator (Rotavapor, Buchi, Switzerland) at 41°C at 150 rpm. CRISPR gRNA (crRNA) were obtained by transcribing template oligonucleotides (Eurofins Genomics, Louisville, KY) using MEGAscript™ T7 Transcription Kit (Invitrogen, Carlsbad, CA, USA) according to the manufacturer’s protocol. crRNA was purified using Oligo Clean & Concentrator™ (Zymo Research, Irvine, CA, USA). CRISPR ribonuclease complex were formed by adding Cpf1 protein (Integrated DNA Technologies, Coralville, IA) and crRNA in 1M Sodium acetate buffer and incubated for 15 minutes at room temperature. The thin film of LNP material was rehydrated with 1.6 ml of 1 M sodium acetate buffer containing CRISPR complex. The solution was then extruded 10 times each through 800-, 400-, and 200-nm Nuclepore Track-Etch Membrane (Whatman, Little Chalfont, UK) filters with the Lipex Biomembrane extruder (Vancouver, British Columbia, Canada).

### Characterization of iLNP and CRISPR LNP (CR-LNP)

The size and zeta potential of the iLNP and CR-LNP were assessed by the dynamic light scattering technique using Zetasizer instrument (Malvern, Westborough, MA) and dip cell (Malvern Westborough, MA). Encapsulation efficiency of CR-LNP was measured by centrifuging 100 μl of LNP mixture through an Ultracel-100K centrifuge filter (Merck Millipore, Tullagreen, Carrigtwohill) at 21,380 x g for 15 min. Filtrate was collected and A260/280 values were measured on a Nanodrop 2000 spectrophotometer (Thermo Fisher Scientific, USA) and matched to a standard curve to calculate the amount of unencapsulated CRISPR complex to determine overall encapsulation efficiency.

### Cell Culture

The 4T1 murine breast cancer cells were obtained from ATCC and cultured in Roswell Park Memorial Institute (RPMI) 1640 media supplemented with 10% FBS, 1% penicillin–streptomycin. Bone marrow-derived macrophages (BMDM) were obtained by isolation from hind leg bone marrow from Balb/c mice (6–8 weeks, females) and incubation with RPMI 1640 with 10% FBS and penicillin-streptomycin supplementation for 7 days. Cells were incubated with 50 ng/mL IFN-γ and 20 ng/mL LPS for *M1* macrophage polarization when needed. For *M2* macrophage polarization, BMDM were incubated with either 50 ng/mL IL-4 and 50 ng/mL M-CSF or in 1:1 ratio of RPMI and filtered 4T1 conditioned media. For 4T1 sphere formation, we incubated 4,000 4T1 cells with Nanoshuttle-PL and incubated with 3D magnetic spheroid drive from bioprinting kit (Greiner BioOne, Austria) for 2 days to allow sphere formation. All cells were maintained within a 37°C humidified incubator with 5% CO_2_.

### Cell viability and LNP uptake study

Macrophages were cultured in 96-well plates and treated with particles with different compositions for 24 hours. At least 6 wells per condition were tested for in vitro studies. Viability was measured after incubation time using CellTiter-Glo assay according to manufacturer’s protocol. In coculture study, macrophages were treated with CR-Ric-LNP at concentration of between 0-7.5 µg/mL of CRISPR ribonuclease within CR-Ric-LNP. 2,000 macrophages were incubated with each 4T1 spheres in low-attachment 96-well plates (Corning, NY). Viability was measured after 24h of treatment using CellTiter-Glo 3D according to manufacturer’s protocol (Promega, WI).

Uptake of LNP was tested in 2 different celltypes: Thp-1, a human monocytic cell line, and in BMDMs. 10^6^ Thp-1 cells were incubated with Rhodamine-labeled CR-LNP and uptake was measured by flow cytometry 2 hours after treatment. To assess uptake mechanisms, CR-LNP uptake was also measured in the presence of various uptake inhibitors. 5,000 BMDMs per well were treated with Chlorpromazine (10 µg/mL), genistein (100µM), cytochalasin D (1µM), or Nocodazole (20µM) (Sigma Aldrich, St. Louis, MO) for 4 hours, before incubation with Rhodamine-labeled CR-LNP for 2 hours. Cells were fixed using 4% paraformaldehyde and counterstained with DAPI for cell detection. Rhodamine signal intensity was quantified on a single-cell basis using automated segmentation within NIS Elements.

LNP uptake was normalized with the mean value of Rhodamine signal from untreated macrophages incubated with CR-LNP.

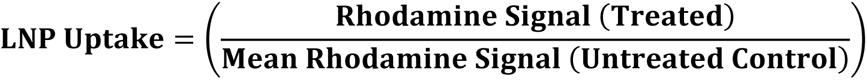

Three independent wells per condition were imaged at 20× magnification, capturing ≥5 fields per well to ensure representative sampling. Data are presented as mean ± SEM, statistical significance was determined via one-way ANOVA with Tukey’s post hoc test (GraphPad Prism v9.0).

### CRISPR gene editing efficacy studies

We conducted proof-of concept study with CRISPR targeting GFP. 10,000 cells of GFP-expressing MDA-MB-463 cells were seeded in 96-well plates and treated with CR-GFP-LNP. GFP signal was visualized using fluorescence microscopy after 3 days of incubation to determine the knock down. Additional studies were conducted to connect uptake and CRISPR editing efficacy in 4T1 and BxPB3 cell lines. 50,000 cells were seeded in 96 well plates, treated with RhodamineB-labeled CR-luc-LNP targeting luciferase knock down. Uptake by 4T1 and BxPC3 cell lines were evaluated at time points 0.5, 1, 2, 4, 6, and 24h by measuring Rhodamine fluorescence signal. Further, knock down of luciferase was measured on cells seeded with the same condition in 96-well white plates for optimized signal measurements. 10µl of Renilla luciferin (200mg/mL) were pipetted to each wells and luciferase activity was measured with Biotek Synergy™ H4 plate reader (Biotek, Winooski, VT) immediately. Double strand break activity was detected in BMDMs seeded in 8-well chamber slide system (Nunc lab-Tek, Waltham, MA).

### CRISPR screening for macrophage modulating genes

We did functional CRISPR screening using our CR-LNP platform evaluating genes that may affect macrophage polarization. crRNA targeting at least 2 sites per genes were designed using benchling.com. CR-LNPs were formed using the crRNAs and macrophage modulating efficiency was evaluated in macrophages incubated with 4T1 conditioned media in 1:1 ratio with RPMI media to mimic TME. Macrophage polarizations were evaluated by CD206 and CD80 staining 1 day after CR-LNP treatment. Cells were stained with a CD80 antibody (Invitrogen, UK) for M1 phenotype, CD206 antibody (Abcam, UK) for M2 phenotype and imaged using fluorescence microscopy. Number of cells expressing CD206 and/or CD80 were quantified and displayed as cell number per field of view. At least 9 field of view were analyzed for each conditions.

Effects of CRISPR treatment to macrophage polarization were also evaluated in RNA level. 100,000 BMDM were seeded in 6-well plates with and without M2 conditioned media (IL-4 and M-CSF). Cells were treated with Sham CR-LNP of CR-Ric-LNP for 24h and total RNA was harvested using RNeasy^®^ Mini Kit (Qiagen, Hilden, Germany). Purified RNA samples were then submitted to Genewiz (South Plainfield, NJ, USA) for RNA sequencing.

To assess the double strand breaks upon CRISPR treatment, BMDMs were treated with CR-Ric-LNP and fixed after different time points between 5 and 360 minutes of incubation using 4% paraformaldehyde. Cells were stained with γ-H2AX antibody for double strand break detection and counterstained with DAPI. Further, we used GeneArt^TM^ Genomic Cleavage Detection kit to detect indels caused by CRISPR treatment.

To evaluate the changes in macrophages treated with CR-Ric-LNP in the presence of tumor cells, we incubated varying number of CR-Ric-LNP treated and untreated macrophages together with 4T1 spheres. Infiltration of macrophages and changes in macrophage marker expression was followed 24h after coculture. Macrophages were stained with CD206 and CD80 and imaged using confocal microscopy to track their localization within 4T1 spheres.

### *In vivo* Model of Breast Cancer Liver Metastasis

Female Balb/c mice, 6 to 8 weeks old, were purchased from Charles River Laboratories. The mice were maintained in animal facilities at Houston Methodist Research Institute approved by the American Association for Accreditation of Laboratory Animal Care and in accordance with current regulations and standards of the United States Department of Agriculture, Department of Health and Human Services, and NIH (Bethesda, MD). All the surgical procedures are approved by the Institutional Animal Care and Use Committee of Houston Methodist Research Institute.

To generate liver metastasis, 10^5^ breast cancer 4T1 cells were injected into the spleen of female Balb/c mice as previously described^113^, followed by splenectomy. The cells injected in the spleen will disseminate to the liver via portal vein and will start forming macro-metastases (metastasis >2mm) around day 16. Mice were monitored daily following their surgery.

### Biodistribution Assessment of CR-LNP Efficacy

Eighteen days after initiation of liver metastasis, animals were randomly distributed into five groups (n=6/group): i) untreated control; ii) CR-LNP 1-hour; iii) CR-LNP 2-hours; iv) CR-LNP 4-hours; v) CR-LNP 24-hours. Animals were imaged with IVIS at the appropriate time points, then harvested. Brain, heart, liver, lungs, and kidneys were isolated, weighed, and excised and weighed to be homogenized for biodistribution analysis. The fluorescence intensity of cell lysates was measured at λ_ex_ = 545 nm and λ_em_ = 560 nm to detect Rhodamine B signal. The obtained values were normalized to the theoretical injection concentration of Rhodamine B-conjugated LNPs. Portions of each organ were frozen in OCT (optimal cutting temperature) after fixation with 4% formalin. Organs were later sectioned on a CryoStar NX50 cryostat (Epredia) at 10 μm for immunofluorescence and IMC analysis.

### Single Cell RNA Sequencing

Female animals were treated with CR-Ric-LNP or PBS at day 18 after 4T1 liver metastasis initiation, 3 animals per group were sacrificed 24h after treatment and liver metastases lesions were harvested for scRNA-seq analysis. Liver metastasis tissues were dissociated, and single cell RNA libraries were constructed using the 10x Genomics Chromium 5’ v2 kit by EMPIRI, Inc (Houston, USA) according to manufacturer’s protocol.

cDNA amplification and fragmentation were performed with adjusted cycles for optimal coverage. Libraries were sequenced on NovaSeq 6000 Illumina platform to a target depth of 50,000 reads/cell.

### Single Cell sequencing data analysis

Raw FASTQ files were aligned to the refdata-cellranger-mm10-1.2.0 mouse reference genome and quantified using Cell Ranger v9.0.1 (10x Genomics).

Single-cell RNA sequencing (scRNA-seq) data were processed using *Seurat* (v5.2.1) in *R* (v4.4.2). Raw count matrices were imported from 10x Genomics .h5 files using the Read10X_h5() function. Cells with fewer than 100 detected genes were excluded. Putative doublets were identified and removed using scDblFinder via the SCP package (v0.5.6), applying an estimated doublet rate of approximately 1% based on sample size. Additional quality control metrics included the number of detected genes (nFeature_RNA), UMI counts per cell (nCount_RNA), the percentage of mitochondrial gene expression (percent.mt), ribosomal RNA content (riboRatio), and library complexity (log10GenesPerUMI). Cells expressing fewer than 200 genes or exhibiting greater than 15% mitochondrial content were excluded from downstream analysis. Quality distributions were visualized using violin and density plots.

Following filtering, expression data were normalized using log normalization with a scaling factor of 10,000. Differential gene expression between experimental groups was computed using the wilcoxauc() function from the *presto* package (v1.0.0), which applies a Wilcoxon rank-sum test and ranks genes by area under the curve (AUC). Top DEGs per cluster were used for manual annotations. Automated annotation was conducted using SingleR with MouseRNAseqData reference dataset for cross-validation.

Gene set enrichment analysis (GSEA) was performed using pre-ranked AUC-based gene lists with the *fgsea* package (v1.3.2), applying a minimum gene set size of 10, and the SCP R package to evaluate the functional relevance of differentially expressed genes.. Mouse gene sets were obtained from the Molecular Signatures Database (MSigDB) using the *msigdbr* package (v10.0.1), specifically the Hallmark (H) collection and KEGG Legacy pathways (C2:CP:KEGG_LEGACY). Significantly enriched pathways were defined as those with a false discovery rate (FDR) adjusted p-value < 0.05. Normalized enrichment scores (NES) were calculated and used to classify pathways as upregulated or downregulated. Pseudotime dynamics were inferred using Monocle3 with myeloid population. Cell-cell communications were analyzed by applying CellChat v1.6.0 to ligand-receptor databases to infer cellular communication crosstalk.

### Immunofluorescence Analysis

Organ sections were mounted onto slides and sections were permeabilized with 1% Triton X-100 then blocked with 10% BSA and 1% FC block (BD Pharmingen) in 1x PBS pH 7.2. Primary and secondary antibodies were added in sequence to slides for 45 minutes at 37°C in a humidified container. Finally, slides were preserved with Prolong Gold antifade reagent (Invitrogen, Eugene, OR) and sealed with coverslip.

Slides were imaged using a Nikon Eclipse Ti inverted microscope with motorized stage and automated z-drive was used for full-slide analysis. Fluorophore excitation/emission filters were optimized to minimize spectral spillover (DAPI, FITC, TRITC, and Cy5). Regions of interest (ROIs) were manually selected to cover the entire tissue section with tile stitching mode with 10% overlap percentage.

Sections of slides that were also identified to contain similar metastases sizes were sent to IMC core at Houston Methodist Hospital for analysis with antibody panels (Table 1). Stitched images from immunofluorescence imaging and IMC were imported into QuPath for cell segmentation and colocalization analysis.

### Image Analysis Using QuPath

Quantitative analysis was performed using QuPath (v0.4.3)^114^, an open-source software for digital pathology. DAPI channel was used to identify and segment cell nuclei using a cell detection algorithm with optimized parameters for size, intensity threshold, and background correction. Nuclei centers were extracted as coordinate points for downstream spatial analysis. Cell boundaries were defined by expanding a fixed radius (e.g., 5–10 µm) from the nuclear centroid to approximate the cellular area. Tumor cells were identified based on specific markers (Pan-CK), and immune cell subtypes (e.g., CD8+/CD4+ T cells, CD11b+/F4/80+ macrophages) were classified accordingly. Immune cell localization relative to tumor cells was categorized into two spatial zones: (i) Immediate neighbor (0–30 µm) includes immune cells within a 30 µm radius of a tumor cell centroid, and (ii) adjacent region (30–100 µm), which includes immune cells located between 30 µm and 100 µm from the nearest tumor cell. Batch effects were minimized by normalizing fluorescence intensities across slides using reference controls.

## Supplementary Figures

**Supplementary Figure 1.**
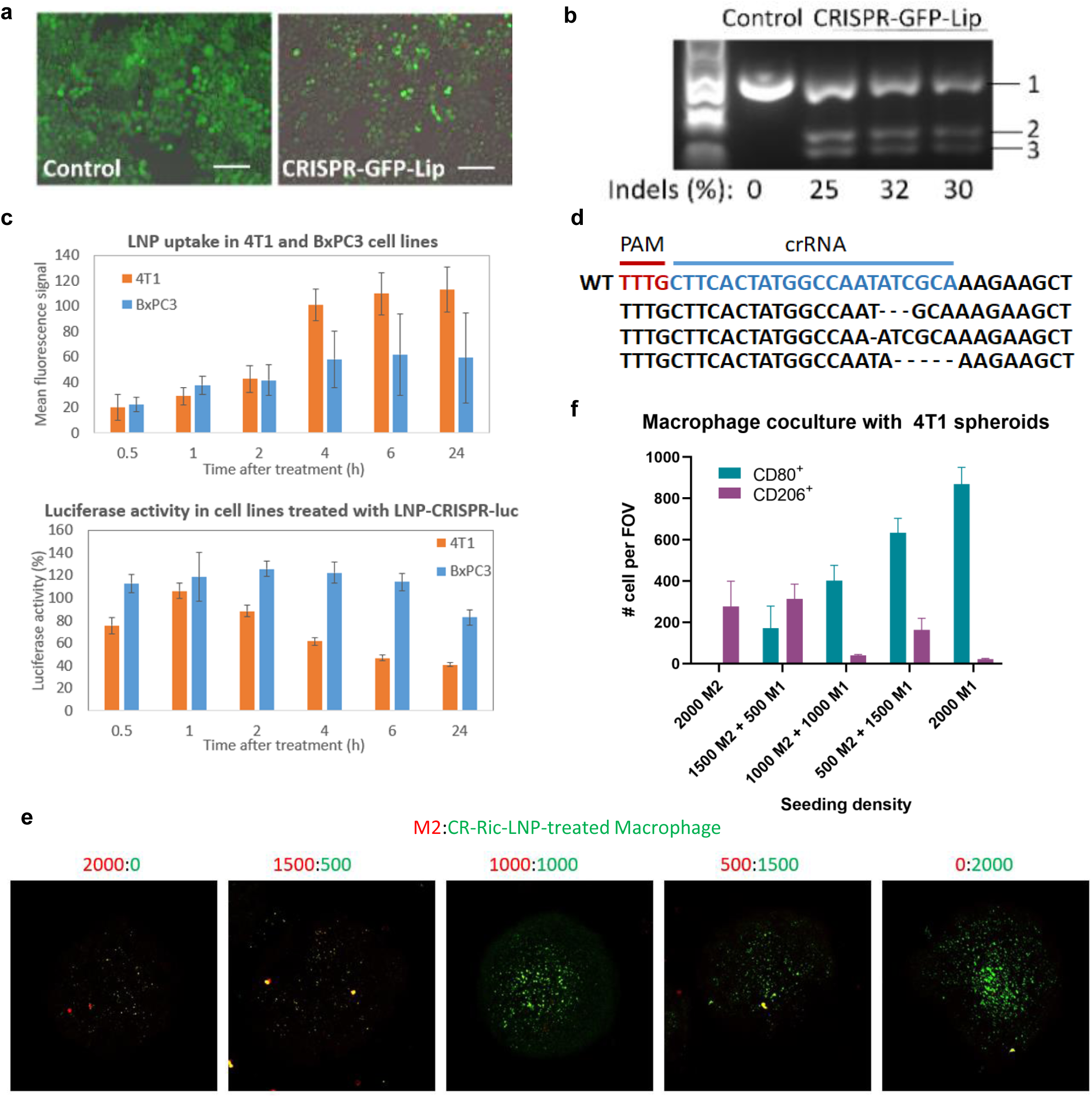
CR-LNP testing and validation. a) Liposome loaded with CRISPR system targeting GFP knocked out GFP expression in MDA-MB-436 cells. B) Gene deletion detected via GeneArt cleavage detection kit, mismatches were recognized & cleaved by detection enzyme, resulting in two distinct bands (band 2&3) from parental band (band 1) in gel electrophoresis (n=3). Scale bar=100 µm. b) Proof of principle testing in breast cancer and pancreatic cancer cell lines indicate that CRISPR activity directly correlates to the uptake by the cell lines. c) Representative gene mutations within Rictor exon in the cells treated with CR-Ric-LNP. e) Macrophage cocultured with 4T1 spheroid models showed recruitment of CR-Ric-LNP-treated (M1) macrophages within the spheres, imaged via fluorescence microscopy. f) Quantification of the macrophage subtypes infiltrating 4T1 tumor spheres

**Supplementary Figure 2.**
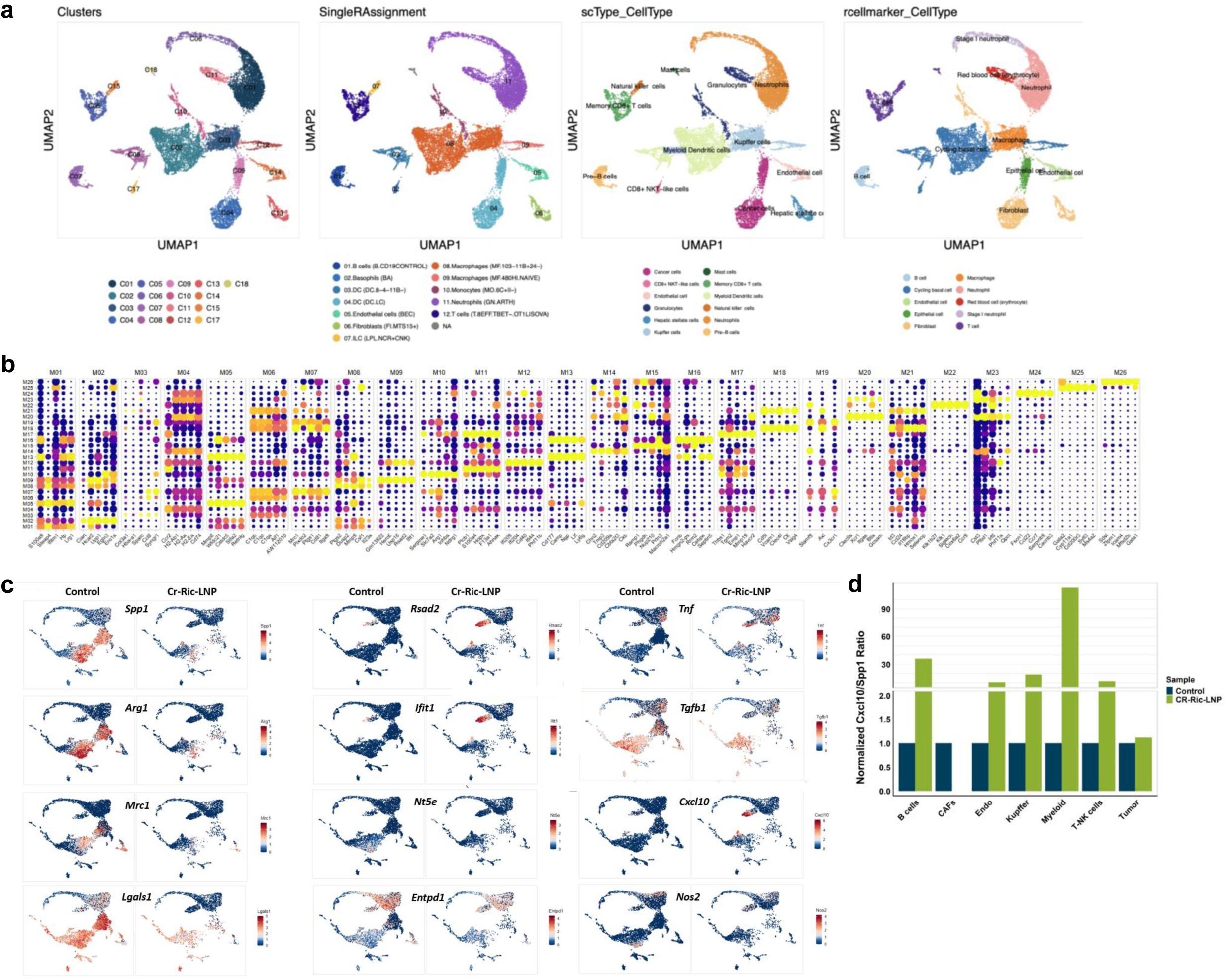
a) Single-cell RNA sequencing (scRNA-seq) data from a breast cancer liver metastasis model were annotated using SingleR with reference transcriptomic datasets to identify major immune, tumor, and stromal cell populations. B) Dot plots display the top 5 most significantly upregulated (yellow) genes across *de novo*-annotated myeloid clusters in the breast cancer liver metastasis model. Dot size indicating the percentage of cells expressing the gene (*expression frequency*) and color intensity reflecting the average log2 fold change (*expression level*). c) Feature plots visualizing the expression of key functional markers across *de novo*-annotated myeloid clusters in a breast cancer liver metastasis model for immunosuppressive signature (e.g., *Spp1, Arg1, Mrc1, Lgals, Tgfb1, Nt5e, Entpd1*), Interferon (IFN)-response signature (e.g., *Ifit1, Rsad2, Cxcl10*), and inflammatory signature (e.g., *Nos2, Tnf*). Each panel overlays gene expression (gradient: blue → red) onto UMAP embeddings. d) The Cxcl10/Spp1 expression ratio was calculated across myeloid subpopulations to quantify their functional polarization in breast cancer liver metastases. Negative values indicate a predominance of *Spp1* (immunosuppressive programs, e.g., pro-tumorigenic macrophages), while positive values (red) reflect *Cxcl10*-dominant inflammatory states (e.g., anti-tumor myeloid activation). Treatment with *Rictor*-targeting CR-LNPs significantly increased the Cxcl10/Spp1 ratio across most myeloid clusters, indicating a broad reduction in immunosuppressive phenotypes and a shift toward immunostimulatory functions.

**Supplementary Figure 3.**
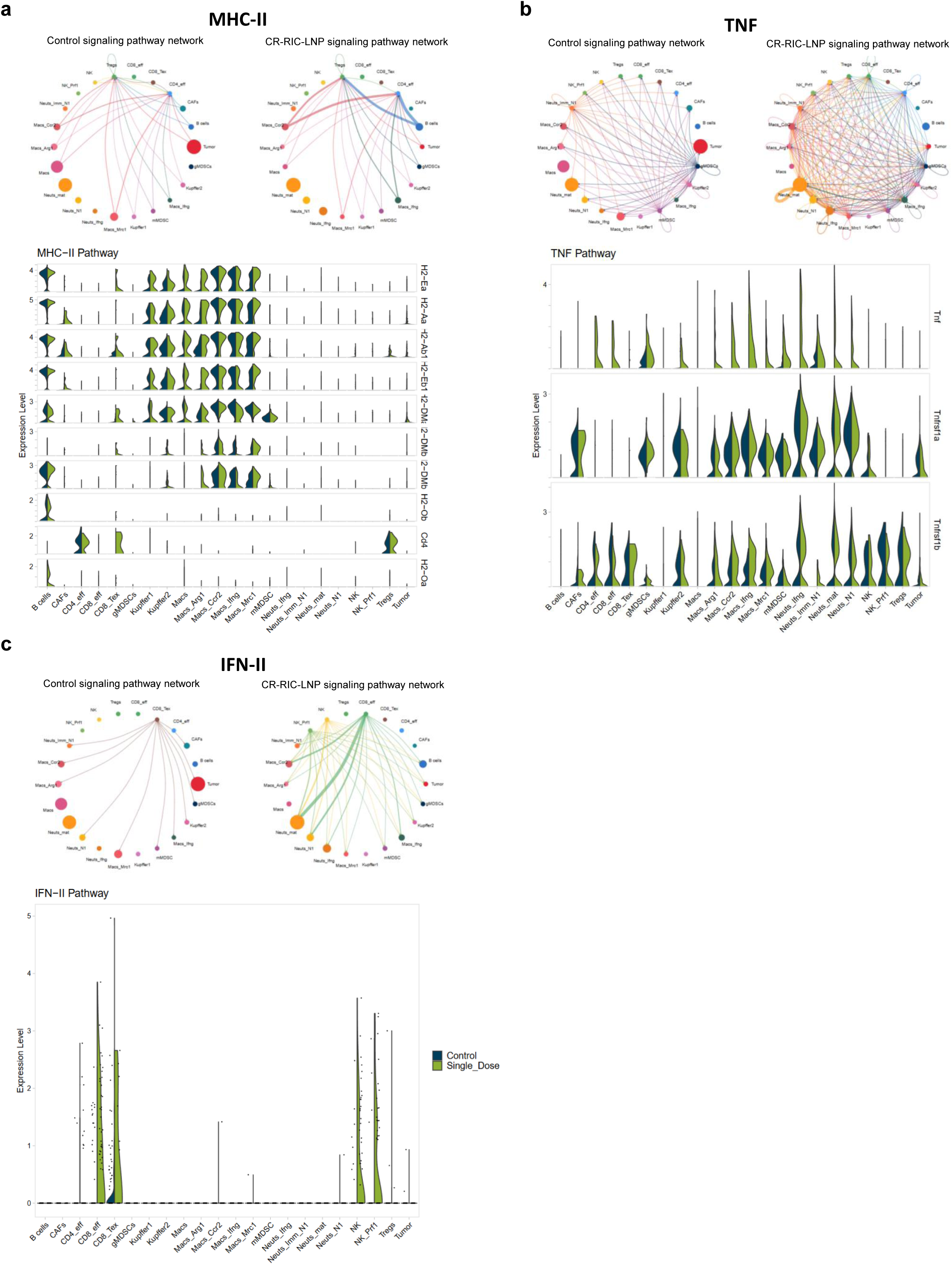
L-R violin plot of gene expression within immunomodulatory pathways. Violin plots display average expression level in each clusters and circular plots display weighted cellular interaction between clusters of a) MHC-II pathway, b) VEGF pathway, and c) TNF pathway. The y-axis represents an expression level of genes in log (TPM).

**Supplementary Figure 4.**
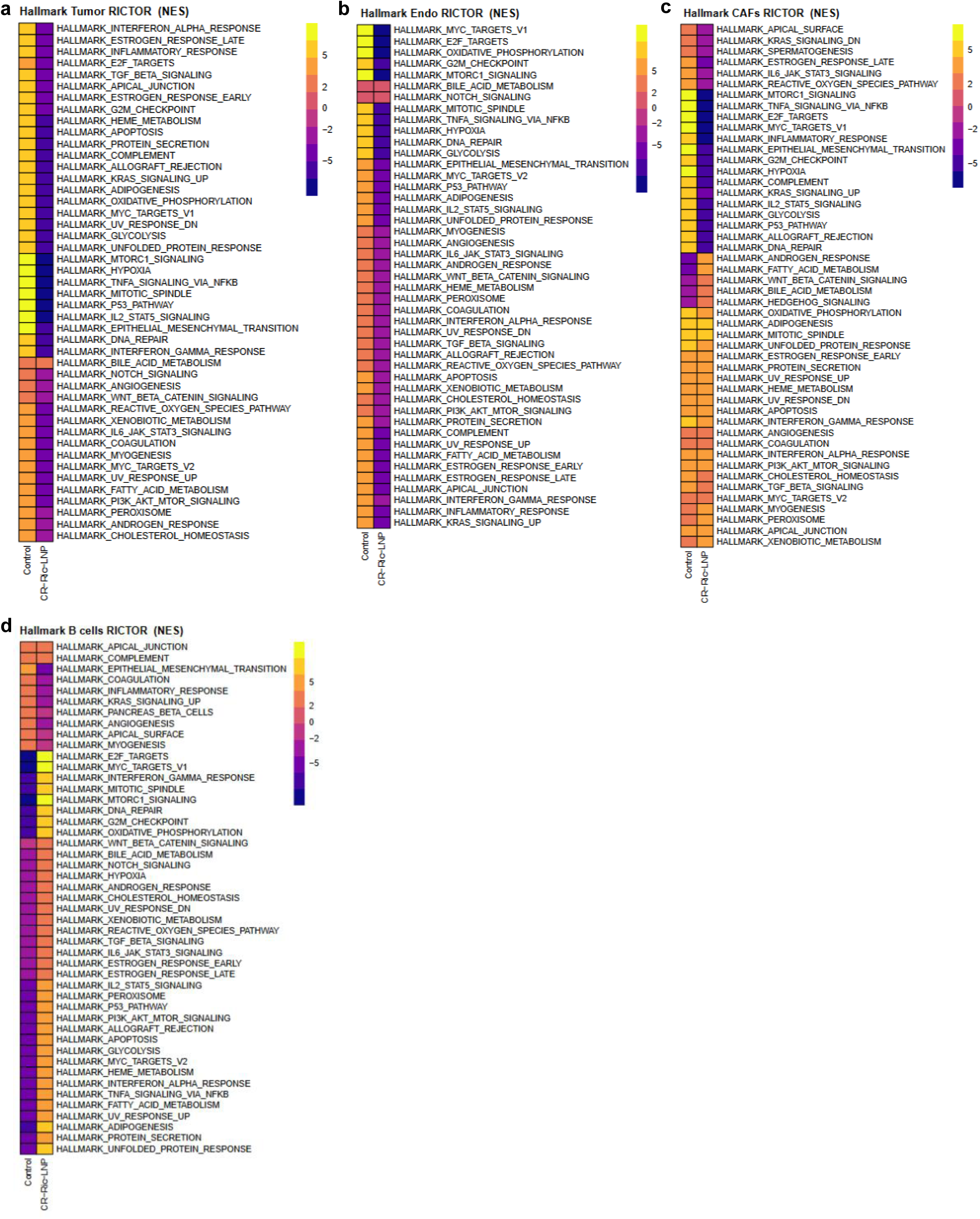
GSEA Hallmark NES analysis of a) tumor, b) Endothelium, c) CAF, and d) B-cell clusters.

**Supplementary Figure 5.**
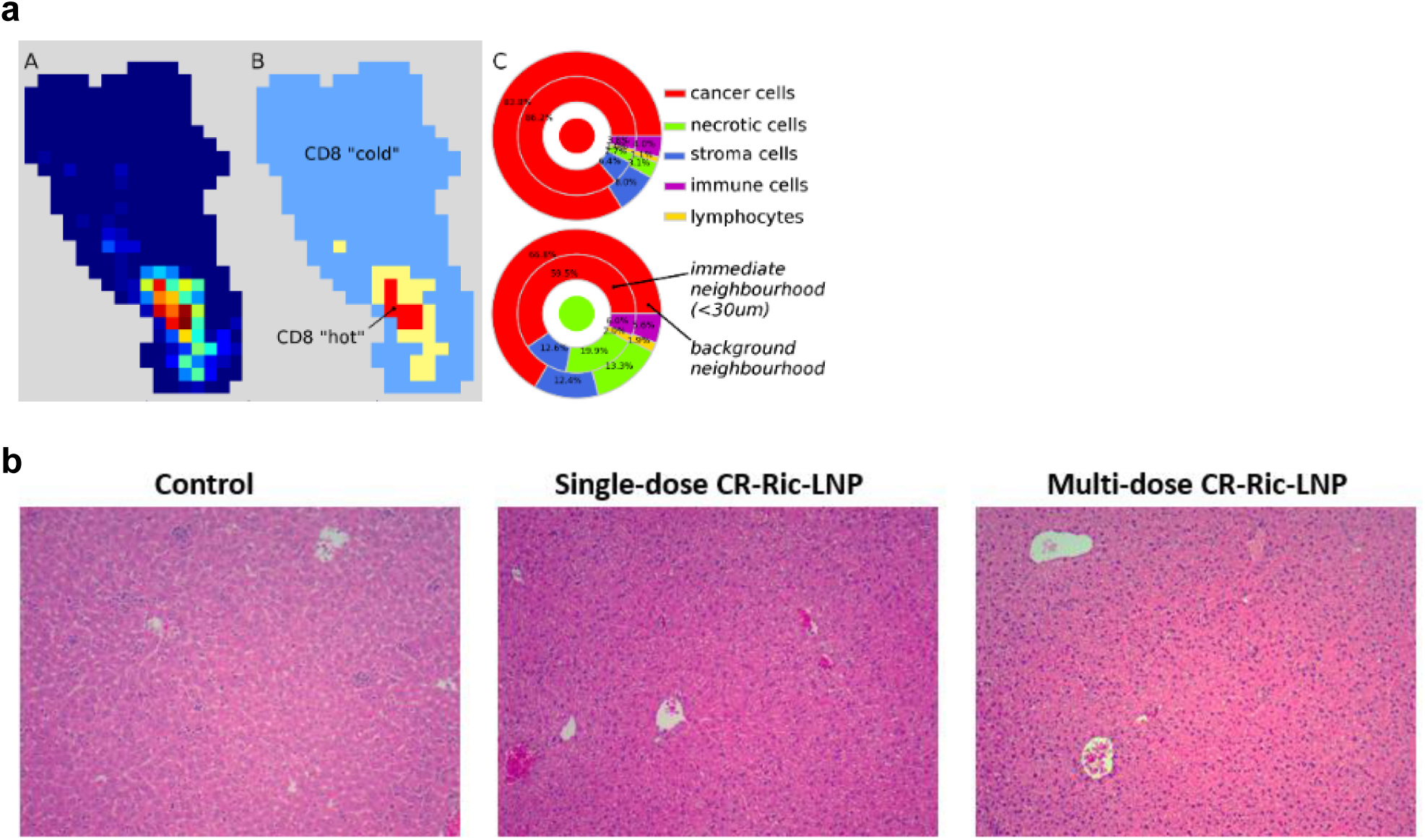
a). Example of cell spatial segmentation and analysis from tumor biopsy of: A) all cell density heatmap showing cell density distribution in tumor (high density; low density); B) tissue segmentation into “hot” and “cold” tissue areas based on local CD8 cell density; C) relative cell distribution showing cellular composition surrounding each cell; cancer and necrotic cell local cellular microenvironment from cancer biopsies. b) Representative H&E staining from uninvolved liver samples after single and multiple dose of CR-Ric-LNP.

